# Random forest modelling of neuropathological features identifies microglial activation as an accurate pathological classifier of C9orf72-related amyotrophic lateral sclerosis

**DOI:** 10.1101/2021.12.10.471808

**Authors:** Olivia M. Rifai, James Longden, Judi O’Shaughnessy, Michael D.E. Sewell, Karina McDade, Michael J.D. Daniels, Sharon Abrahams, Siddharthan Chandran, Barry McColl, Christopher R. Sibley, Jenna M. Gregory

**Affiliations:** Translational Neuroscience PhD Programme, Centre for Clinical Brain Sciences, University of Edinburgh, 49 Little France Crescent, Edinburgh EH16 4SB; Centre for Clinical Brain Sciences, University of Edinburgh, 49 Little France Crescent, Edinburgh EH16 4SB; UK Dementia Research Institute, University of Edinburgh, 49 Little France Crescent, Edinburgh EH16 4SB; Euan MacDonald Centre for Motor Neurone Disease Research, University of Edinburgh, 49 Little France Crescent, Edinburgh EH16 4SB; Centre for Discovery Brain Sciences, University of Edinburgh, 15 George Square, Edinburgh EH8 9XD; Human Cognitive Neuroscience-Psychology, School of Philosophy, Psychology and Language Sciences, University of Edinburgh, 7 George Square, Edinburgh EH8 9JZ; Institute of Quantitative Biology, Biochemistry and Biotechnology, School of Biological Sciences, University of Edinburgh, The King’s Buildings, Edinburgh EH9 3FF; Simons Initiative for the Developing Brain, University of Edinburgh, 15 George Square, Edinburgh EH8 9XD

**Author notes:** Corresponding authors: Olivia M. Rifai, Jenna M. Gregory, Fax: No fax available.

**Keywords:** Amyotrophic lateral sclerosis, frontotemporal dementia, C9orf72, neuroinflammation, microglia, post-mortem tissue

## Abstract

Amyotrophic lateral sclerosis (ALS) and frontotemporal dementia (FTD) are regarded as two ends of a pathogenetic spectrum, termed ALS-frontotemporal spectrum disorder (ALS-FTSD). However, it is currently difficult to predict where on the spectrum an individual will lie, especially for patients with *C9orf72* hexanucleotide repeat expansions (HRE), a mutation associated with both ALS and FTD. It has been shown that both inflammation and protein misfolding influence aspects of ALS and ALS-FTSD disease pathogenesis, such as the manifestation or severity of motor or cognitive symptoms. Previous studies have highlighted markers which may influence *C9orf72*-associated disease presentation in a targeted fashion, though there has yet to be a systematic and quantitative assessment of common immunohistochemical markers to investigate the significance of these pathways in an unbiased manner. Here we report the first extensive digital pathological assessment with random forest modelling of pathological markers often used in neuropathology practice. This study profiles glial activation and protein misfolding in a cohort of deeply clinically profiled *post-mortem* tissue from patients with a *C9orf72* HRE, who either met the criteria for a diagnosis of ALS or ALS-FTSD. We show that microglial immunohistochemical staining features, both morphological and spatial, are the best independent classifiers of disease status and that clinicopathological associations exist between microglial activation status and cognitive dysfunction in ALS-FTSD patients with *C9orf72* HRE. Furthermore, we show that spatially resolved changes in FUS staining are also an accurate predictor of disease status, implying that liquid-liquid phase shift of this aggregation-prone RNA-binding protein may be important in ALS caused by a *C9orf72* HRE. Our findings provide further support to the hypothesis of dysfunctional immune regulation and proteostasis in the pathogenesis of *C9orf72* ALS and provide a framework for digital analysis of commonly used neuropathological stains as a tool to enrich our understanding of clinicopathological associations between cohorts.

## Introduction

Amyotrophic lateral sclerosis (ALS) is a clinically heterogeneous neurodegenerative disease, with substantial variation in prognosis between patients and disease duration ranging from months to decades[61]. The most common known monogenetic cause of ALS is a hexanucleotide (G_4_C_2_) repeat expansion (HRE) in the first intron of the *C9orf72* gene, accounting for approximately 25% of familial and 10% of sporadic cases (C9-ALS)[20, 36, 54]. *C9orf72* HREs are associated with diverse clinical presentations, ranging from ALS and ALS-frontotemporal spectrum disorder (ALS-FTSD)[51], to parkinsonism and psychosis[2, 9, 21, 40, 49, 63]. Clinical trials in people with ALS are often based on phenotypic outcome measures like survival, functional rating scores (FRS) such as the ALS-FRS, and cognitive scores such as the Edinburgh Cognitive and Behavioural ALS Screen (ECAS). Thus, the clinical heterogeneity seen in C9-ALS, even with the same underlying monogenetic cause, is a hugely complicating factor in identifying potential therapeutics. Understanding the molecular drivers of clinical heterogeneity will enable more targeted and personalised therapies to be explored and improve stratification for clinical trials aimed at reducing specific symptom burden.

Thus far, pathological examination of *post-mortem* tissue from over 200 cases with a *C9orf72* HRE exists in the literature, primarily in ALS and FTD cases[12–17]. This collection of research suggests that, despite differences in clinical presentation, *C9orf72* mutations are very similar pathologically, exhibiting unifying features such as a p62-positive granule cell layer[4, 25, 57], dipeptide repeat (DPR) protein deposits[6, 19, 35], and cortical aggregates of the RNA-binding protein (RBP) TDP-43[25, 40, 49, 57]. They are further differentiated from sporadic ALS pathology by TDP-43-negative, p62-positive staining[25]. Indeed, other RBPs are seen to misfold certain ALS subtypes, but whether they are involved in C9-ALS remains to be determined. Interestingly, it has recently been shown that the wildtype form the RBP fused in sarcoma (FUS) mislocalises in sporadic ALS *post-mortem tissue* and vasolisin-containing protein (VCP)-related ALS induced pluripotent stem cell-derived motor neurons; previously this was only documented a property of only the ALS-associated mutant form [59]. Wildtype FUS dysregulation in a *post-mortem* C9-ALS context has yet to be observed, though research demonstrating the association of C9orf72 protein with FUS in stress granules, as well as the negative effects of *C9orf72*-related DPRs on the phase transition regulation of RBPs such as FUS and TDP-43 suggests it is possible[14, 41]. Few studies have shown markers that may be associated with the clinical heterogeneity observed. For example, greater TDP-43 burden in the spinal cord has been previously associated with a shorter disease duration[13]. Additionally, we have previously shown that cognitive impairment is linked with the presence of TDP-43 pathology in extra-motor brain regions, though a subgroup of cases was shown to also have extra-motor TDP-43 pathology despite being cognitively unaffected. Thus, extra-motor TDP-43 pathology is a specific, but not sensitive, marker of cognitive impairment in ALS[26]. While informative, these findings do not account for all clinical heterogeneity observed in C9-ALS. Further investigation into sensitive and specific markers of C9-ALS and associated symptoms using clinically annotated tissue is thus warranted.

Another pathway of interest related to *C9orf72*, and which may influence phenotypic heterogeneity, is immune signalling. Studies examining both *C9orf72* knockdown and haploinsufficiency resulting from HREs have demonstrated immune dysregulation [5, 56]. Additionally, it has recently been suggested that differential inflammatory profiles may influence the manifestation or severity of motor and cognitive symptoms across the spectrum[23]. Indeed, a recent study observed differences in neuroinflammatory profiles in the cerebrospinal fluid of participants with ALS, FTD, or ALS-FTD, though each group possessed similar mutations[45]. Furthermore, levels of inflammatory factors, such as dysfunctional regulatory T cells and levels of interleukins (ILs) in plasma, as well as microglial activation in the cervical corticospinal tract, have been reported to correlate with disease progression[8, 10, 34]. Despite these associations, clinical trials exploring the use of anti-inflammatory drugs for ALS therapy have thus far been unsuccessful[47]. This may be attributed to non-specific mechanisms of action, such as a general blocking of inflammation, when a more nuanced, pathway-specific manipulation may be necessary. Additionally, the bias of preclinical studies toward *SOD1* models must also be considered. Indeed, *SOD1* mutations only account for 2% of ALS cases[53, 54]. As for FTD, there is little focus on anti-inflammatory therapies at present[58].

Previous studies have highlighted some markers which may influence *C9orf72*-associated disease, though there has yet to be a systematic and quantitative assessment of immunohistochemical markers to investigate the significance of these pathways in an unbiased manner. Here we present the first extensive digital pathological assessment of morphological and pathological markers commonly used in neuropathology practice to profile glia-related inflammation and protein misfolding in a cohort of deeply clinically profiled *post-mortem* tissue from patients with a *C9orf72* HRE, who either met the criteria for a diagnosis of ALS or ALS-FTSD. Furthermore, we used machine learning to train a random forest disease classifier using tens of millions of features derived from our digital analysis, to identify relevant markers of C9-ALS status.

## Methods

### Case identification and cognitive profiling

Tissue from 10 ALS *post-mortem* cases with a *C9orf72* HRE was obtained from the Medical Research Council (MRC) Edinburgh Brain Bank (Table 1). All cases had corresponding whole genome sequencing and pathogenic *C9orf72* HRE were confirmed by repeat-primed polymerase chain reaction (PCR). Tissue from 10 age- and sex-matched control cases was obtained from the Edinburgh Sudden Death Brain Bank. Control cases had no history of neurological conditions or evidence of neurodegenerative disease pathology. All *post-mortem* tissue was collected with ethics approval from East of Scotland Research Ethics Service (16/ES/0084) in line with the Human Tissue (Scotland) Act (2006). Use of *post-mortem* tissue for studies was reviewed and approved by the Edinburgh Brain Bank ethics committee and the Academic and Clinical Central Office for Research and Development (ACCORD) medical research ethics committee (AMREC). Clinical data were collected as part of the Scottish Motor Neurone Disease Register (SMNDR) and Care Audit Research and Evaluation for Motor Neurone Disease (CARE-MND)[33] platform, with ethics approval from Scotland A Research Ethics Committee (10/MRE00/78 and 15/SS/0216). Donor patients underwent neuropsychological testing with the Edinburgh Cognitive and Behavioural ALS Screen (ECAS)[1] and all patients consented to the use of their data. Clinical correlates of motor dysfunction/disease progression include disease duration (months) and sequential ALS functional rating score (ALSFRS) data points. Clinical correlates of cognition include Edinburgh Cognitive and Behavioural ALS Screen (ECAS) scores for ALS-specific and ALS non-specific subdomain scores.

**Table 1.** C9orf72 cohort demographics.

|  | Age at death (y) | Disease duration (m) | Region of onset | Cognitive data | ALSFRS |
| --- | --- | --- | --- | --- | --- |
| **Case 1** | 63 | 25 | Lower Limb | Yes (unimpaired) | No |
| **Case 2** | 50 | 29 | bulbar | Yes (language dysfunction) | Yes (4 data points) |
| **Case 3** | 63 | 30 | Upper limb | No | Yes (3 data points) |
| **Case 4** | 62 | 37 | Mixed | Yes (unimpaired) | Yes (6 data points) |
| **Case 5** | 65 | 44 | Lower limb | No | Yes (2 data points) |
| **Case 6** | 65 | 50 | Upper limb | No | Yes (1 data point) |
| **Case 7** | 62 | 50 | Lower limb | Yes (executive dysfunction) | Yes (2 data points) |
| **Case 8** | 43 | 57 | Bulbar | Yes (language dysfunction) | No |
| **Case 9** | 58 | 87 | Lower limb | Yes (language dysfunction) | Yes (3 data points) |
| **Case 10** | 63 | 119 | Upper limb | Yes (FTD – 3 domains) | No |

### Immunohistochemistry (IHC)

Brain tissue was taken *post-mortem* from standardised Brodmann areas (BA) BA4, BA39, BA44, BA46 and fixed in 10% formalin for a minimum of 72 h. Tissue was dehydrated in an ascending alcohol series (70–100%), followed by three 4 h washes in xylene. Three successive 5 h paraffin wax embedding stages were performed followed by cooling and sectioning of formalin-fixed, paraffin-embedded (FFPE) tissue on a Leica microtome into 4 μm thick serial sections that were collected on Superfrost (ThermoFisher Scientific) microscope slides. Sections were dried overnight at 40°C before immunostaining. Sections were dewaxed using successive xylene washes, followed by alcohol hydration and treatment with picric acid for quenching of lipofuscin. For pTDP-43, CD68, Iba1 and FUS staining, antigen retrieval was carried out in citric acid buffer (pH 6) in a pressure cooker for 30 min, after which immunostaining was performed using the Novolink Polymer detection system with a 2B Scientific (Oxfordshire, UK) anti-phospho(409–410)-TDP-43 antibody at a 1 in 4000 dilution, Agilent (Cheadle, UK) anti-CD68 antibody at a 1 in 100 dilution, Abcam (Cambridge, UK) anti-Iba1 antibody at a 1 in 3000 dilution, and Bio-Techne (Minneapolis, USA) anti-FUS antibody at a 1 in 500 dilution. For GFAP staining, no antigen retrieval step was carried out, and immunostaining was performed using the Novolink Polymer detection system with a Dako anti-GFAP antibody at a 1 in 800 dilution. Counterstaining was performed using 3,3’- diaminobenzidine (DAB) chromogen counterstained with haematoxylin, according to standard operating procedures.

### Imaging and digital pathology analysis

Whole tissue sections were scanned with Brightfield at 40x magnification using a Hamamtsu NanoZoomer XR. Using NDP.view2 viewing software (Hamamatsu), regions of interest (ROIs) were taken from key regions for quantification. For both grey and white matter, 3 vascular-adjacent and 3 non-vascular-adjacent regions of interest (ROIs) (500 μm x 500 μm) were taken per tissue section for each stain. ROIs were defined as vascular-adjacent when centered on a vessel larger than a small capillary, with larger, cross-sectional vessels captured where possible. ROIs were considered nonvascular adjacent if they were at least ∼250 μm away from any vessel larger than a small capillary (Figure 1a, Figure S1a). ROIs were analysed using QuPath software[7] superpixel segmentation in a manner previously described[39], with the modification of using difference of gaussians (DoG), rather than simple linear iterative clustering (SLIC) segmentation, to obtain shape features. For vascular-adjacent ROIs, annotations were drawn manually around the vessel (endothelial cell border) and around the area of tissue retraction if present; vessel annotations were subtracted from the image for subsequent automated analyses. Images were segmented into superpixels with stain-specific segmentation parameters selected to best identify the staining of interest (downsample = 1, gaussian sigma = 8.0 for Iba1 and GFAP, gaussian sigma = 5.0 for CD68, FUS and TDP-43). Superpixels were classified using three thresholds for DAB Mean Intensity, according to the pathological precedence of Allred scoring of oestrogen receptor immunohistochemistry in breast cancer[22], and intensity and shape features were calculated for each superpixel. For vascular-adjacent ROIs, distance to vessel (normalised to the square root of the non-vessel-containing area of the image to account for differences in vessel calibre) and distance to superpixels by threshold class were also calculated. Measurements were exported at the image (number of positive detections per threshold class) and superpixel level (threshold class, intensity, shape, and distance features).

**Figure 1.**
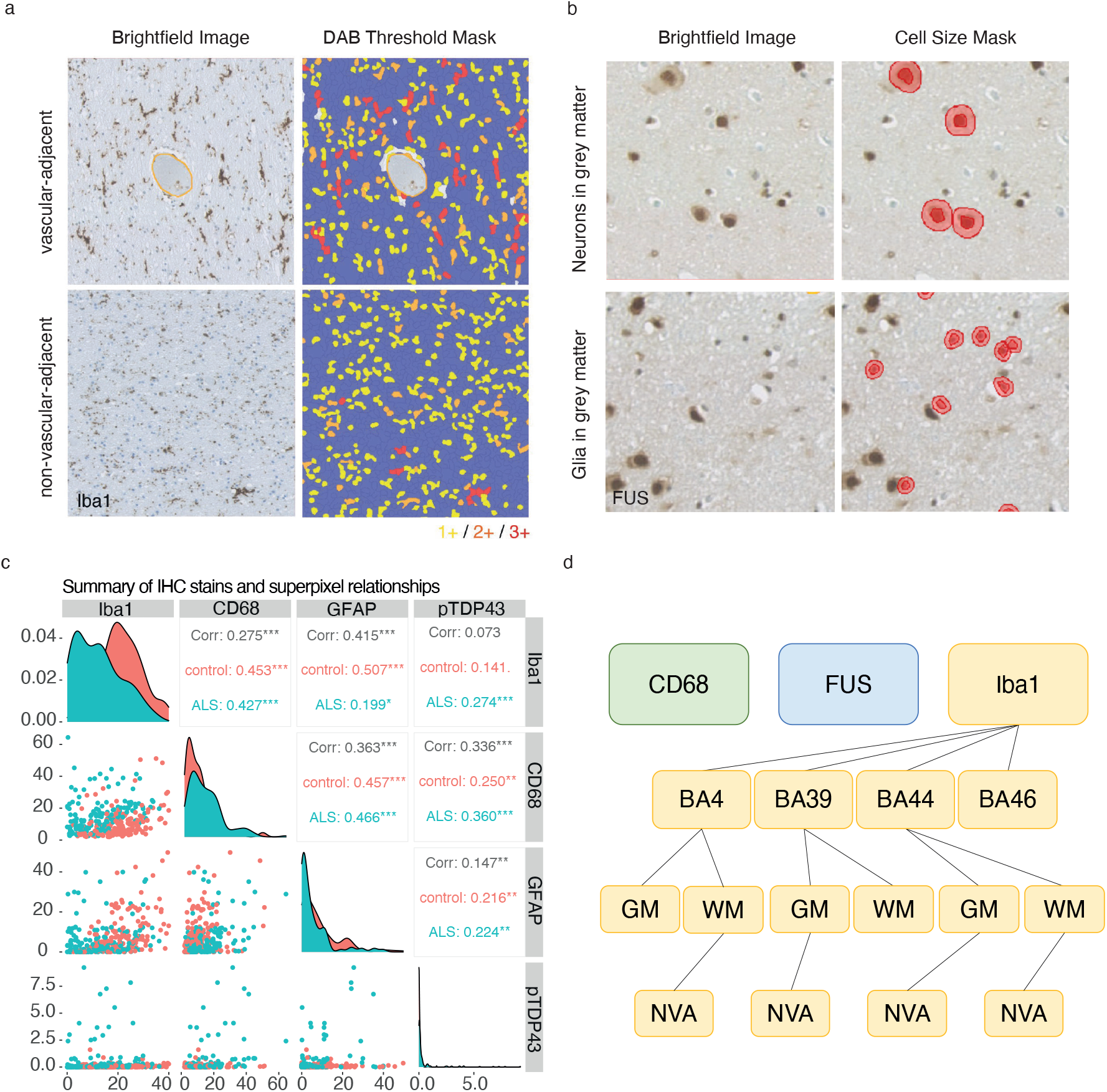
Microglial immunohistochemical staining features are the best independent classifiers of disease status. (a) Example region of interest comprising brightfield images of Iba1-stained tissue in vascular adjacent and non-vascular adjacent regions (left). Iba1 = brown DAB staining with a haematoxylin counterstain. Images on the right show a segmentation-analysis mask overlay created using QuPath digital pathology software coding DAB staining intensity from 1+ to 3+. (b) Example neuronal and glial cell masks created by nuclear area thresholding created using QuPath. (c) Correlation matrix demonstrating inter-relationship between superpixel analysis of histological stains. (d) Hierarchical organisation of results from Random Forest analysis demonstrating accuracy of each stain as a disease classifier. Note Iba1 demonstrates multi-level accuracy (i) stain, (ii) brain region, (iii) sub-region 1 (white matter vs grey matter) and (iv) sub-region 2 (vascular-adjacent and non-adjacent). BA, Brodmann area; GM, grey matter; WM, white matter; NVA, non-vascular-adjacent; VA, vascular-adjacent. Spearman, * p < 0.05, ** p < 0.01, *** p < 0.001, **** p < 0.0001.

Nuclear and cytoplasmic FUS localisation were quantified using cell segmentation in QuPath. For neuronal quantification, only grey matter ROIs were included, cells were segmented (pixel size = 0.5 μm, background radius = 0, median filter radius = 0, sigma = 4, minimum nuclear area = 150 μm^2^, maximum nuclear area = 5000 μm^2^, intensity threshold = 0.1, max background intensity = 0, cell expansion = 10 μm) and nuclear, cytoplasmic, and whole cell intensity and shape features were calculated and exported. For glial quantification, both grey and white matter ROIs were included, cells were segmented (pixel size = 0.5 μm, background radius = 0, median filter radius = 0, sigma = 4, minimum nuclear area = 10 μm^2^, maximum nuclear area = 100 μm^2^, intensity threshold = 0.1, max background intensity = 0, cell expansion = 5 μm) and nuclear, cytoplasmic, and whole cell intensity and shape features were calculated and exported. Scripts used in these analyses can be found in the Supplementary Information. Manual TDP-43 grading was performed as described previously[26] and was visualised using GraphPad Prism (version 9.2.0).

Data were visualised in RStudio (R version 4.1.1) with “ggplot2” (version 3.3.5) and “Ggally” (version 2.2.1) packages. Datasets that were found to be normally distributed via Shapiro-Wilk’s test were subjected to parametric tests; unpaired t-test for two-group comparisons, and ANOVA with Tukey’s multiple comparisons correction for three-group comparisons. Datasets that were found to be non-normal via Shapiro-Wilk’s test were subsequently subjected to non-parametric tests; Mann-Whitney U for two-group comparisons, pairwise Wilcoxon with Holm-Sidak multiple comparisons correction for three-group comparisons and Spearman’s test for correlations. Statistical comparisons were only conducted between groups with n ≥ 3. Results were averaged over 3 ROIs capturing the same region and presented as ungrouped or grouped by brain region, grey or white matter, and vascular or non-vascular adjacent. For box plots, data were further averaged when certain stratifications were not included to avoid pseudoreplication (e.g., when comparisons were made between grey and white matter within a brain region without grey matter vascular-adjacent and grey matter non-vascular-adjacent subregions, vascular-adjacent and non-vascular adjacent results were averaged for grey matter and white matter). For correlations, all stratified data were included even when data were ungrouped to explore relationships between two stains across all images. Multiple linear regression models were fitted where significant correlations were found, and coefficients were compared by t-tests on individual regression coefficients.

### Machine learning

The automated analysis of IHC images in QuPath generated approximately 80.2 million superpixel features and 5.2 million cell segmentation features, 56.4 million of which (66%) were used to train random forest classifiers using disease versus control datasets. Classifiers were trained with 1000 trees per classifier and Information Gain as the split criterion. Classifiers were then tested on the remaining 33% of the dataset. Cross-tables were generated for regions and X^2^ tests were performed to determine which regions allowed the random forest to correctly predicted disease significantly better than chance. Sensitivity (100*true positives/total disease) and specificity (100*true negatives/total control) scores were also calculated (Supplementary Table 1). Finally, leave-one-out analyses were performed to determine the predictive power of each feature (Supplementary Table 2).

### RT-qPCR

Reverse transcription quantitative polymerase chain reaction (RT-qPCR) was conducted by extracting RNA from human FFPE using the RNAstorm FFPE RNA extraction kit (Cell Data Sciences) on two 10 μm curls per sample cut from BA4. RNA was eluted in 50 μL nuclease-free water, after which sample concentrations were measured using a NanoDrop 1000 (ThermoFisher) and concentrated further for 10-15 minutes at 45°C using an Eppendorf Concentrator Plus if needed. 125 ng of RNA was reverse transcribed using Superscript IV (Life Technologies) as per the manufacturer’s instructions, after which 10 ng cDNA was amplified with PowerUp SYBR Green Master Mix (ThermoFisher) and forward and reverse primers to *FUS* and housekeeping 18S ribosomal RNA gene *18Sr* (0.5 μM), selected to control for age- and degradation-related artefact. FUS primers were specifically designed to target two non-contiguous central portions of the sequence so as to better detect product that may be partially degraded due to the nature of FFPE tissue preparation and storage[3]. Primer sequences are listed below, with alternate *FUS* primer sequences that failed to amplify cDNA listed in Supplementary Information.

*18Sr* Forward: 5’-GGCCCTGTAATTGGAATGAGTC-3’

*18Sr* Reverse: 5’-CCAAGATCCAACTACGAGCTT-3’

*FUS* Forward: 5’-ACTTCCTTTCTTTTGCTCTCAC-3’

*FUS* Reverse: 5’-ATTAAGCCCCACTAAGCCC-3’

Quantitative PCR was run on eight control and eight C9-ALS samples in 10 ng triplicates using qTOWER^3^ 84 (Analytik Jena GMBH) (50°C for 2 min, 95°C for 2 min, 40 × 95°C for 5s, 60°C for 30 s). One control was removed due to off-target amplification as determined by a melting curve. Cycle threshold (Ct) values for *FUS* were normalised (ΔCt) to *18Sr* Ct values, and fold changes (2^-ΔΔCt^) were calculated relative to the average ΔCt of control samples. Six controls and seven cases that did not exhibit off-target amplification and were consistent among triplicates were included in the final analysis. Conditions were visually inspected for homogeneity of variance and an unpaired t-test was performed to test significance between conditions.

## Results

### Microglial immunohistochemical staining features are the best independent classifiers of C9-ALS

To investigate pathological differences between C9-ALS and controls, *post-mortem* tissue from a deeply clinically phenotyped cohort (Table 1) was stained with antibodies commonly used in the neuropathological work-up of ALS cases in all brain banks. Specifically, we examined antibody immunoreactivity to microglia-enriched marker Iba1, microglia/macrophage lysosomal activation marker CD68, astrocyte activation marker GFAP, and RNA-binding proteins FUS and TDP-43, which are known to mislocalise and aggregate in ALS[44, 59]. Threshold-based stain intensities, as well as morphology and spatial features, were calculated using superpixel and cell segmentation methods in QuPath (Figure 1a-b) for the ALS-vulnerable motor cortex (BA4) and for three vulnerable cognitive regions related to the domains assessed in the ECAS[1]; BA39 (language), BA44 (fluency) and BA46 (executive). These were further stratified by non-vascular-adjacent and vascular-adjacent regions in grey and white matter. Collectively this equated to 4,800 images for digital analysis, with 240 images per case. To determine the relationship between the total number of stain-positive superpixels between stains and identify differences between C9-ALS and control cases, correlation analyses were performed. Significant positive correlations were found between the number of Iba1+ and CD68+ superpixels in disease (R = 0.427 p < 0.001) and controls (R = 0.453, p < 0.001) (Figure 1c). Additionally, significant positive correlations were found between the number of CD68+ and pTDP-43+ superpixels in disease (R = 0.360, p < 0.001) and controls (R = 0.250, p < 0.01) (Figure 1c). The number of GFAP+ superpixels was found to positively correlate with CD68+ superpixels in disease (R = 0.466, p < 0.001) and controls (R = 0.457, p < 0.001), demonstrating consistency between glial activation markers (Figure 1c). Notably, this did not simply represent an increase in cell number; see digital cell recognition analysis (Figure S1b-c). Finally, total GFAP+ superpixels were also found to positively correlate with pTDP-43+ superpixels in C9-ALS (R = 0.224, p < 0.01) and controls (R = 0.216, p < 0.01), suggesting a relationship between astrogliosis and TDP-43 aggregation (Figure 1c). When visually inspected, the data spread was most distinct between C9-ALS and controls for Iba1+ versus CD68+ and CD68+ versus pTDP43+ correlations, the details of which are further characterised in Figure 3.

Classification potential of immunohistochemical staining features was interrogated using a random forest machine learning model to determine whether the model could be trained to predict C9-ALS versus control status based on phenotypic tissue features. Overall, digital analysis of immunohistochemical staining conducted in this way was found to be an accurate disease classifier with 69% sensitivity (X^2^ p < 0.0001) and 66% specificity (X^2^ p < 0.0001) overall (Figure 1d; Supplementary Table 1). Interestingly, all regions were independently predictive of C9-ALS status, irrespective of whether the regions were clinically involved (i.e., motor versus cognitive involvement (Supplementary Table 1). When broken down by stain, CD68, FUS and Iba1 stains were both sensitive (i.e., 67%, 64% and 78%, X^2^ p < 0.0001, p < 0.001, p < 0.0001, respectively) and specific (i.e., 60%, 69% and 84%, X^2^ p < 0.05, p < 0.0001, p < 0.0001, respectively) when predicting disease status, while GFAP staining was sensitive (79%, X^2^ p < 0.0001) but not specific (53%, X^2^ p = 0.527), and pTDP-43 staining was specific (64%, X^2^ p < 0.001) but not sensitive (51%, X^2^ p = 0.8744) to disease status, consistent with previous findings (Supplementary Table 1) [27]. This meant that GFAP and pTDP-43 staining yielded more false positive and false negative results, respectively. This may imply that the astrocyte activation marker used in this study is not an adequate predictor of C9-ALS status, though this does not rule out the potential usefulness of other astrocyte-related markers that were not tested, such as ALDH1L1 and AQP4. FUS and CD68 staining were accurate classifiers of disease overall, though this was not maintained within subregions. Iba1 was the most predictive stain, with accurate classification in all four brain regions, in most grey and white matter regions within those and even at the vascular level in BA39 non-vascular-adjacent grey and white matter. The predictive power of microglia-related stains underscores disease-related changes to microglia in C9-ALS and reveals microglia-related stains to be sensitive and specific classifiers of disease status. Finally, almost all included morphological and spatial features had some predictive power for classifying disease. Notably, the features with the highest predictive power for disease included neuronal and glial Haematoxylin and DAB parameters from FUS cell segmentation, distance of negative superpixels to vessel, percentage of class 1+ superpixel detections, and solidity and circularity of class 2+ superpixel detections (Supplementary Table 2).

Interestingly, FUS localisation features from our cell segmentation analysis appeared to have more predictive power of disease status than superpixel count features (Supplementary Table 2) which may also act as a proxy measure for neuronal loss, as FUS is a predominantly nuclear protein. However, no significant reductions in the number of neurons, glia, or superpixels detected in FUS analyses were found between C9-ALS and controls (Figure S1b-d); in fact, a significant increase in glia in BA44 white matter and superpixels detected in BA44 and BA46 white matter were detected in C9-ALS (Figure S1b-d). Given previous reports of FUS mislocalisation and nuclear and cytoplasmic condensates in sporadic ALS[59], and the involvement of C9orf72 protein in FUS-related stress granule dynamics[14], we next sought to assess for digitally apparent differences in localisation and aggregation of FUS in our cohort. Further investigation of nuclear and cytoplasmic FUS staining differences revealed significant increases in nuclear/cytoplasmic mean intensity ratio in BA4 and BA46 glia, but not in neurons (Figure 2a-c). We next interrogated whether this change in intensity was associated with increased FUS expression, or rather nuclear FUS aggregation, using reverse transcription quantitative polymerase chain reaction (RT-qPCR). *FUS* primers were specifically designed to target a central region of the sequence so as to better detect product that may be partially degraded due to the nature of FFPE tissue preparation and storage[3], and revealed no significant difference in *FUS* expression between C9-ALS and controls (Figure 2d). Further support to the notion that this effect could be related to failed degradation and aggregation of FUS, rather than expression differences, is that cytoplasmic condensates in both neurons and glia are readily identifiable in these cases (Figure 2e).

**Figure 2.**
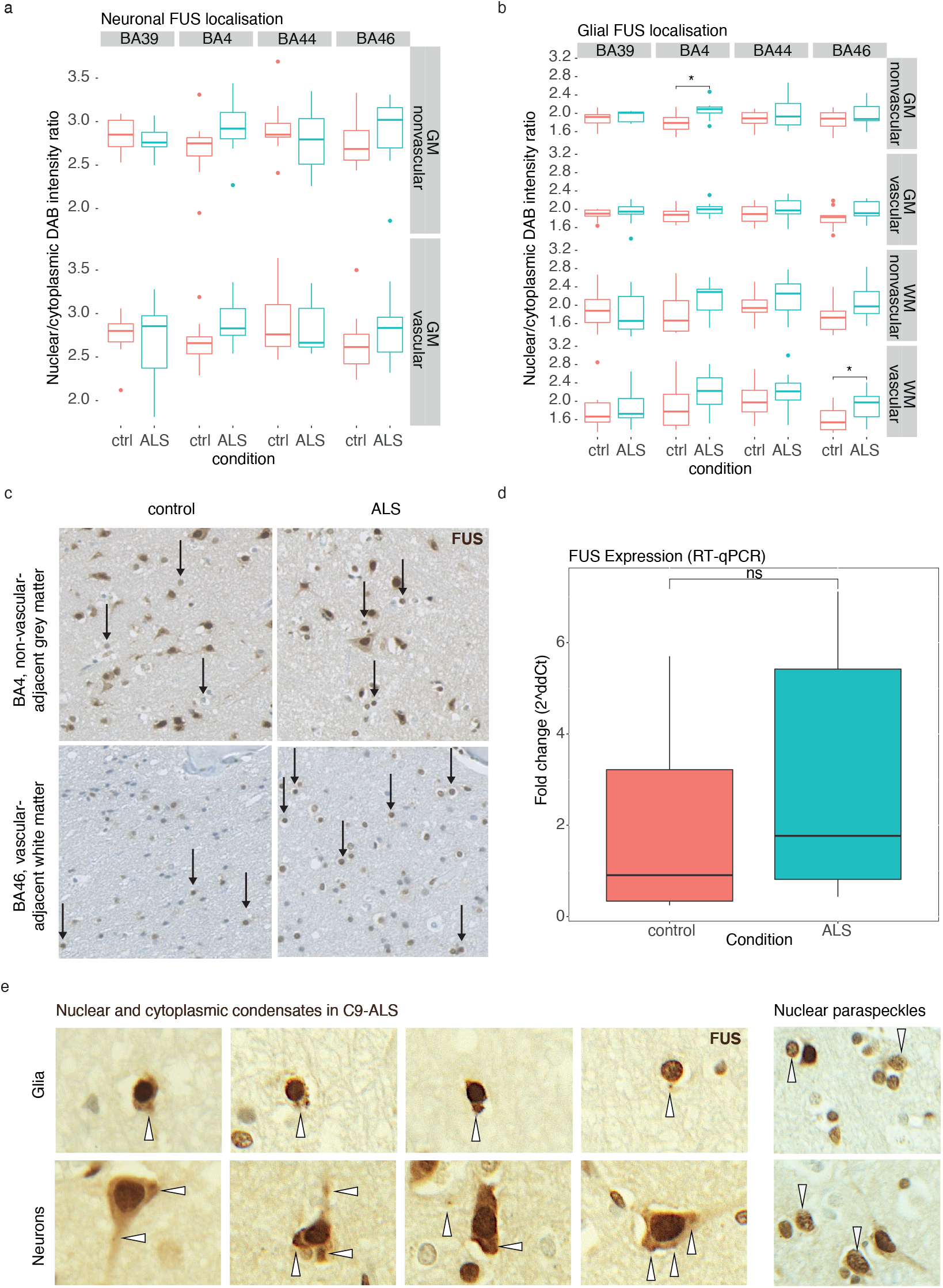
Glia exhibit changes in FUS localisation. Box and whisker plots demonstrating (a) neuronal (unpaired t-test) and (b) glial (Mann-Whitney U test) nuclear/cytoplasmic ratio of FUS immunostaining intensity by brain region, demonstrating increased nuclear/cytoplasmic ratio in disease. (c) Representative images of glial FUS staining with significant differences between control and ALS cases. (d) RT-qPCR of FUS expression demonstrating no significant difference between control and disease (unpaired t-test). (e) Example FUS nuclear and cytoplasmic condensates and nuclear paraspeckles in C9-ALS cases. BA, Brodmann area; GM, grey matter; WM, white matter; NVA, non-vascular-adjacent; VA, vascular-adjacent. * p < 0.05, ** p < 0.01, *** p < 0.001, **** p < 0.0001.

### C9-ALS post-mortem tissue exhibits microgliosis-related changes

As features of the microglia-related stains Iba1 and CD68 were the most accurate predictors of C9-ALS status, and distribution of total positive superpixels for these stains between C9-ALS and controls appeared to be the most distinct, we further explored these stains and their relationships at a regional level. Investigation of the aforementioned microglia-related correlations with multiple linear regression revealed that y-intercepts (disease: y = 10.5437, control: y = 1.9610), but not slopes (disease: a = 0.3733, control: a = 0.4162), of the linear models of Iba1- versus CD68-positive superpixels were significantly different between C9-ALS and control cases (p < 0.001), indicating elevated microgliosis in disease (Figure 3a). This general pattern is consistent across most regions, and although power is reduced when the data are stratified, significant correlations persist in BA4 white matter regions (Figure 3b). For the relationship between CD68- and pTDP-43-positive superpixel number, multiple linear regression revealed that both slopes (disease: a = 0.0528, control: a = 0.001497; p < 0.001) and y-intercepts (disease: y = -0.24604, control: y = 0.065258; p < 0.01) of linear models were significantly different between C9-ALS and controls (Figure 3c), suggesting a relationship between microglial activation and TDP-43 aggregation in disease. A general relationship between CD68 and pTDP-43 is preserved particularly across grey matter regions, as well as BA4 and BA39 white matter (Figure 3d). However, the only significant correlation was found in BA4 vascular-adjacent grey matter, which may be due to the low number of pTDP-43+ detections combined with low power resulting from regional stratification.

**Figure 3.**
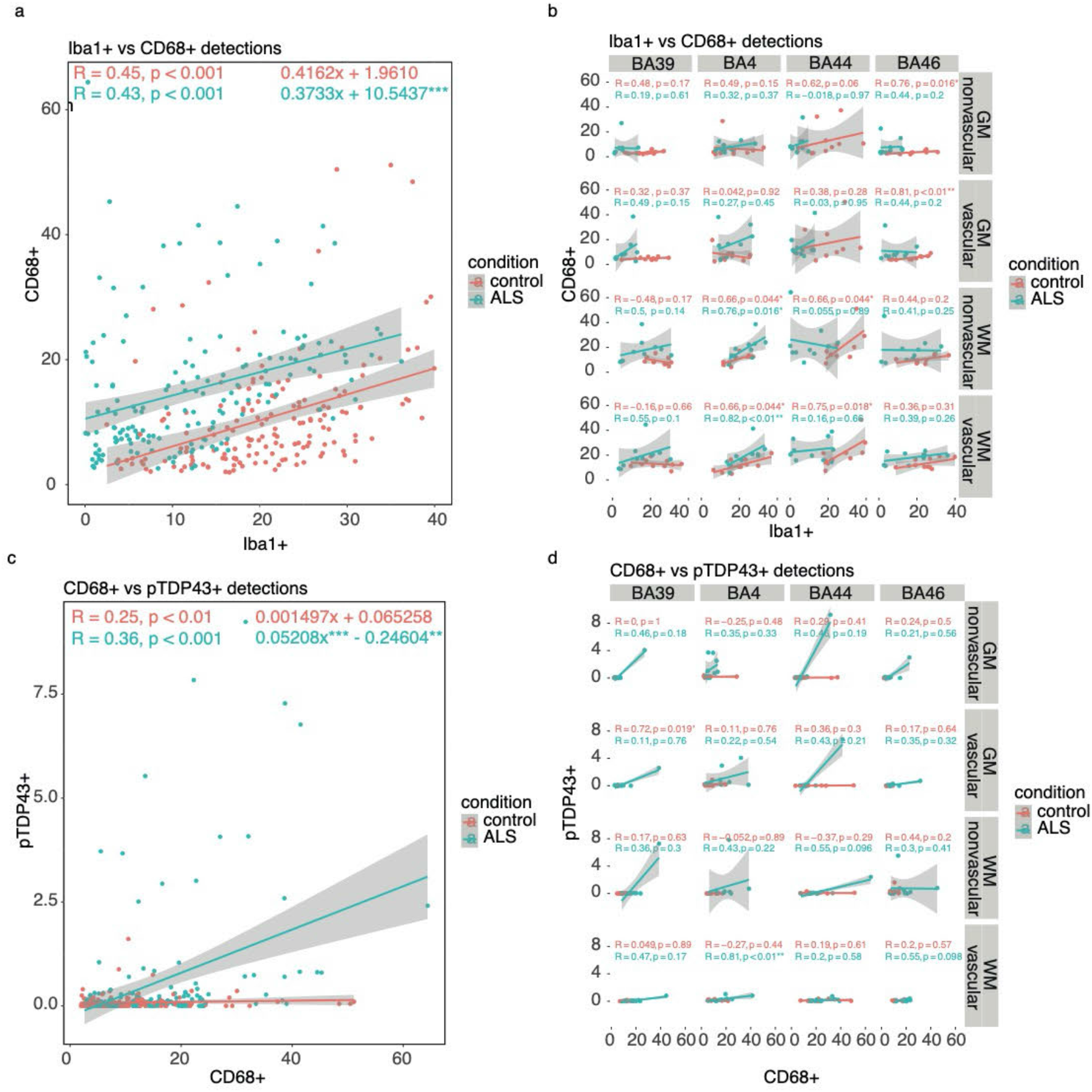
C9-ALS post-mortem tissue exhibits microgliosis-related changes. (a) Scatter plot demonstrating the relationship between CD68 (activated microglial stain) and Iba1 (total microglial stain) demonstrating an overall increase in activated microglia in disease vs control. (b) Correlation matrix plotting data from (a) by brain region subdivided in to grey and white matter and vascular adjacent and non-adjacent regions. (c) Scatter plot demonstrating the relationship between CD68 (activated microglial stain) and pTDP-43 staining demonstrating an increase in activated microglia with increasing TDP-43 staining. (d) Correlation matrix plotting data from (c) by brain region subdivided in to grey and white matter and vascular adjacent and non-adjacent regions. BA, Brodmann area; GM, grey matter; WM, white matter. Spearman, * p < 0.05, ** p < 0.01, *** p < 0.001, **** p < 0.0001.

Analysis of specific features of the strongest classifier of C9-ALS status, Iba1, revealed significant increases in Iba1-positive superpixel area in all included white matter regions as well as in extramotor grey matter (Figure 4a). Interestingly, there were also significant decreases in superpixel circularity of class 1+ and class 3+ Iba1 superpixels (digital mask demonstrated in Figure 1a) between disease and control cases, predominantly in white matter regions (Figure 4b). While this might be unexpected, as microglial activation morphology is more circular and ameboid compared to the ramified homeostatic morphology[55], it must be noted that these findings reveal changes to area and circularity of superpixels containing Iba1 stain, thus indicating disease-related morphological changes of a non-physiological nature and are not necessarily suggestive of activation status. Finally, significant decreases in the distance of class 1+ and class 2+ Iba1 superpixels from the central vessel were observed in C9- ALS, predominantly in BA4 grey matter (Figure 4c), which may reflect migration to vessels as a response to systemic inflammation or compromised blood-brain barrier integrity[29]. It is also possible that the combined result of morphological changes and increased vessel-adjacent Iba1+ staining is indicative of an increased presence of non-microglial, Iba1+ monocyte-derived cells that have migrated from vessels in disease (Figure 4c).

**Figure 4.**
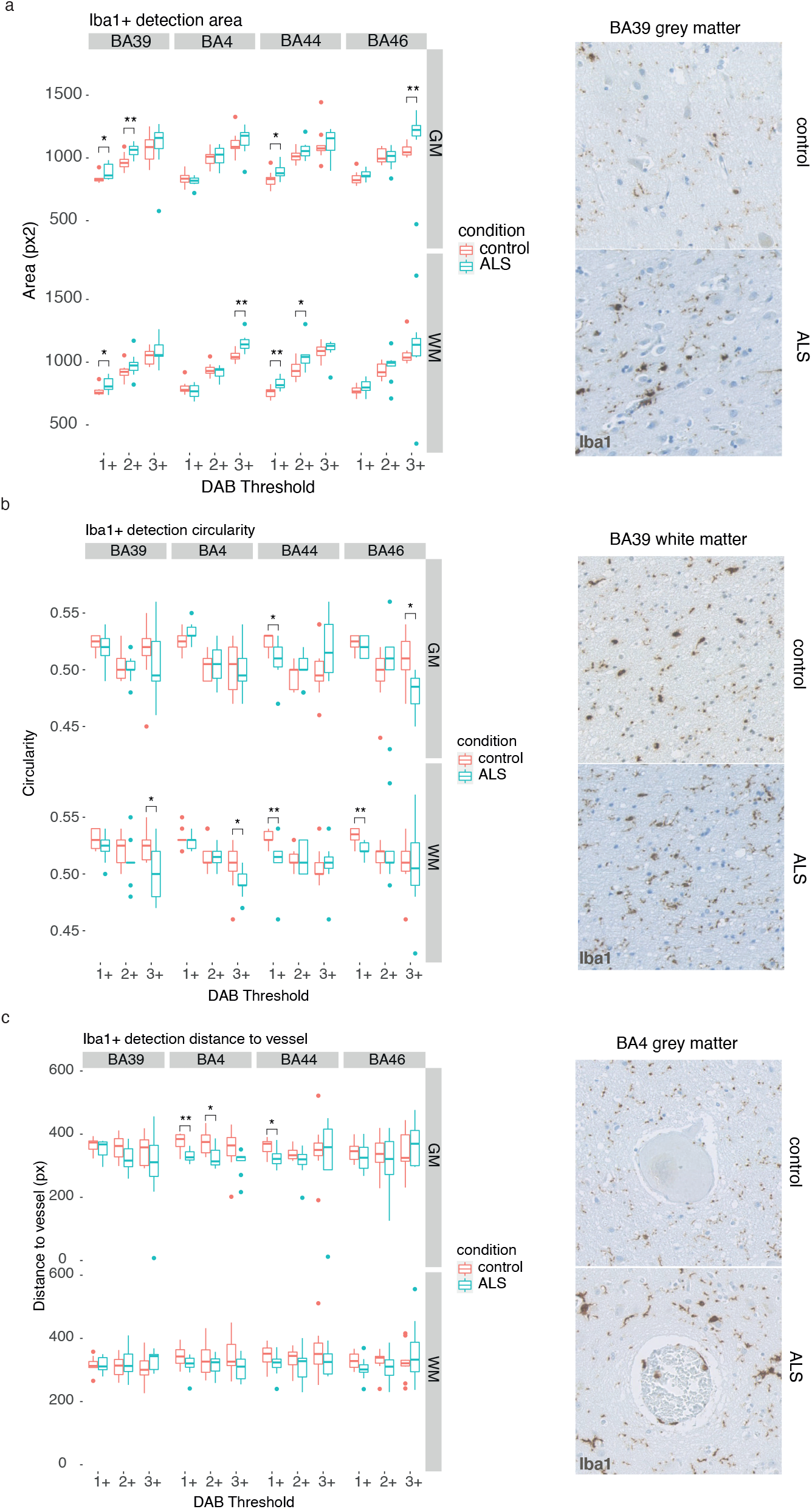
Analysis of microglial staining features reveals morphological and spatial differences in C9-ALS-FTD. (a-c) Examples of feature analysis for Iba1 staining, demonstrating differences in staining intensity (a), morphology (b) and proximity to blood vessels (c) in disease vs control cases. Example photomicrographs on right show control (above) and disease (below). Intensity classes 1-3 represent increasing DAB intensity, with 1 = mild, 2 = moderate, 3 = high. BA, Brodmann area; GM, grey matter; WM, white matter. Mann-Whitney U test, * p < 0.05, ** p < 0.01, *** p < 0.001, **** p < 0.0001.

### Clinicopathological associations exist between microglial activation status and cognitive dysfunction

Findings from immunohistochemistry data were next analysed for relationships with clinical features by comparing cognitively unimpaired C9-ALS cases to those with C9-ALS-FTSD. The amount of CD68+ and pTDP- 43+ staining in relevant regions was assessed between language-impaired cases as determined by ECAS. Language-impaired ALS cases (n = 4) were found to have significantly higher CD68+ staining in BA39 grey matter compared to controls (n = 10) in both grey (p < 0.05) and white (p < 0.01) matter, while this was not the case for language-unimpaired cases (n = 3) (Figure 5a-c). Both C9-ALS subgroups (language-impaired and language-unimpaired) had more CD68+ superpixels per Iba1+ superpixel number in BA39, suggestive of elevated microglial lysosomal activation in ALS cases as mentioned previously (Figure 3), with significant differences in y-intercepts of linear models (language-impaired: y = 7.482, p < 0.001; language-unimpaired: y. = 4.1073, p < 0.05) compared to controls (y = 3.878) (Figure 5c). Interestingly, the slope of the linear model for language-impaired cases was significantly increased compared to controls (impaired: a = 1.037, control: a = 0.172, p < 0.05), while this was not the case for language-unimpaired cases (a = 0.7689), potentially suggesting a more reactive CD68+ microglial phenotype in language-impaired cases (Figure 5c). Two cases exhibited executive impairment and on average displayed higher CD68+ staining in BA46 grey and white matter, though no statistical analysis was conducted due to low power (Figure S2a). The sole case that did exhibit fluency impairment (C9- ALS-FTD) displayed increased CD68+ staining in BA44 grey and white matter compared to controls and other cases, though, again, no statistical analysis was performed due to absence of replicate cases for verification (Figure S2b). Furthermore, there were no significant differences in pTDP43+ staining in BA39 detected between language-impaired, language-unimpaired and control conditions using the digital analysis method (Figure 5d). When investigating disease duration-related differences, no significant differences in CD68+ staining were found between long survivors (i.e., disease duration of over 48 months[61]), short survivors and controls (Figure 5e). Total pTDP-43+ staining was significantly increased in long survivors in BA4 grey matter (p < 0.05), which may indicate a build-up of aggregates over time (Figure 5f). Finally, no significant differences in total GFAP+ staining were detected in BA39 between language-impaired, language-unimpaired and control cases, or in BA4 between long survivors, short survivors and controls (Figure S2c-d).

**Figure 5.**
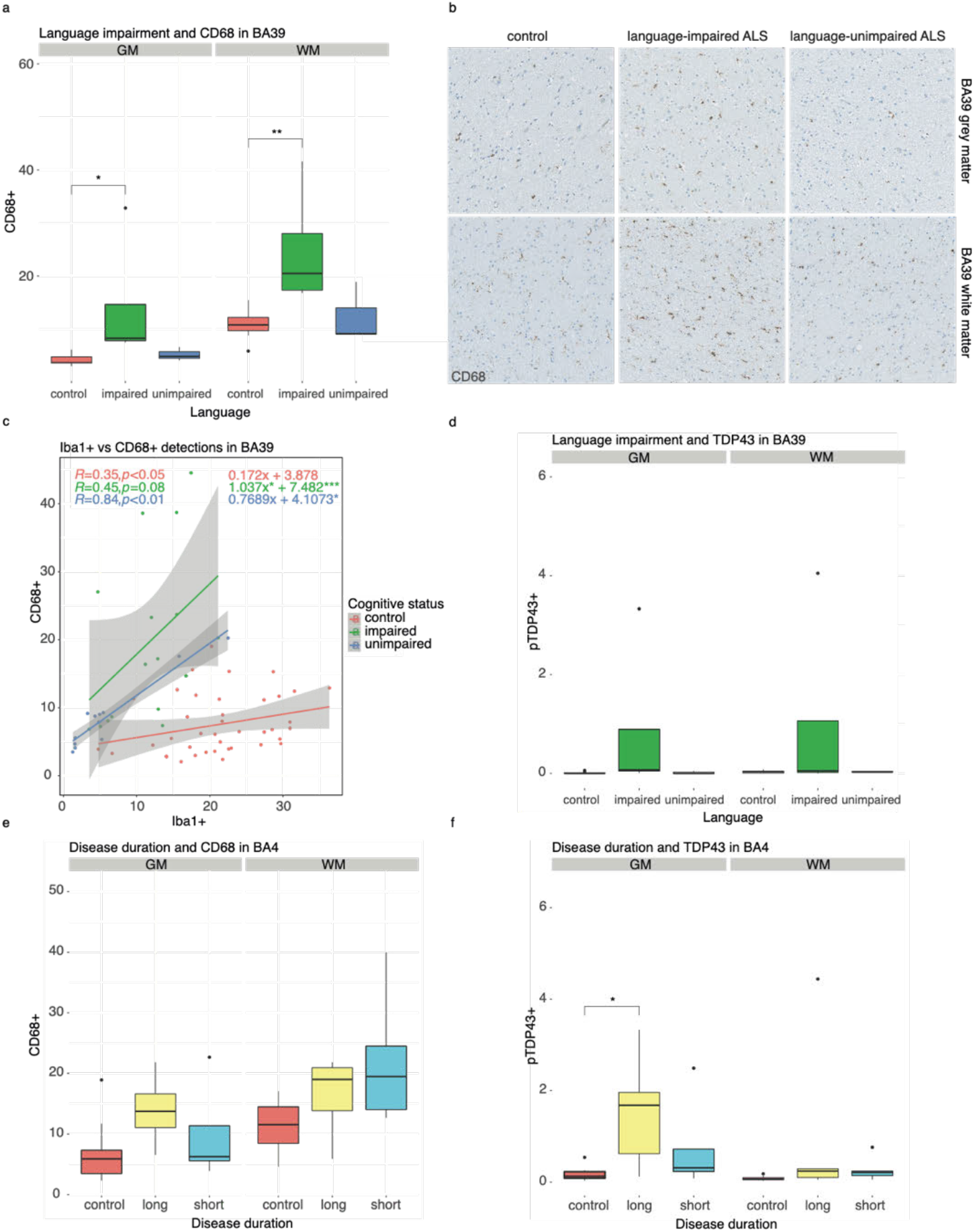
Clinicopathological associations exist between microglial activation status and cognitive dysfunction. (a) Box and whisker plot showing CD68 immunoreactivity in cases grouped by language impairment (controls, language impaired on Edinburgh Cognitive and Behavioural ALS Screen (ECAS) during life and language unimpaired), demonstrating increased immunoreactivity for CD68 in language impaired cases in BA39 – a brain region responsible for language function. (b) Example images (grey matter above and white matter below) with CD68 immunostaining (brown/DAB) and haematoxylin counterstain showing increased immunostaining in language impaired cases in BA39. (c) Scatter plot demonstrating higher CD68 activation status in BA39 of cases vs controls irrespective of cognitive status. (d) Box and whisker plot showing pTDP-43 immunoreactivity in cases grouped by language impairment (controls, language impaired on ECAS during life and language unimpaired), demonstrating increased immunoreactivity for pTDP-43 in language impaired cases in the grey matter of BA39 – a brain region responsible for language function. (e) Box and whisker plot showing CD68 immunoreactivity in BA4 (motor cortex) of cases grouped by disease severity (disease duration from symptom onset; controls, long disease duration and short disease duration). (f) Box and whisker plot showing pTDP-43 immunoreactivity in BA4 (motor cortex) of cases grouped by disease severity (disease duration from symptom onset; controls, long disease duration and short disease duration), demonstrating increased immunoreactivity for pTDP-43 in the grey matter of long survivors in the grey matter. Data in box and whisker plots are averaged across quantified regions not included in the presented stratification, such that each category contains n = number of cases, to avoid pseudoreplication. BA, Brodmann area; GM, grey matter; WM, white matter. Spearman, Wilcoxon with Holm Sidak, * p < 0.05, ** p < 0.01, *** p < 0.001, **** p < 0.0001.

Due to the nature of the automated image analysis and ROI selection, we postulated that milder and sparser TDP- 43 pathology could be underrepresented in our analysis. To test this, TDP-43 aggregate burden was manually graded as described previously[26] (Table 2). As suspected, in BA4, which contained the highest TDP-43 burden across cases, significant differences between neuronal and glial TDP-43 burden in C9-ALS versus control cases were picked up both by the automated and manual methods, though neither method resulted in a significant difference between long and short survivors (Figure S3a). In BA39, which had milder TDP-43 burden overall, the manual method picked up on significant differences between disease and controls that were not detected using the automated method (Figure S3b). However, both methods did not reveal a significant difference between language-impaired and language-unimpaired cases. Thus, while our automated method may be susceptible to underrepresentation of milder pTDP-43 pathology, both methods were similar in their ability to discern differences, or lack thereof, between pTDP-43 pathology in C9-ALS clinical phenotypes. When comparing the scale of differences between language-impaired and language-unimpaired cases for both manually graded TDP- 43 burden and digitally quantified CD68+ staining, there appeared to be a more distinct difference in the latter (Figure S3c). This is likely due to the existence of “mismatch” cases, or ALS cases that exhibit TDP-43 burden in cognitive regions but did not experience related cognitive impairment[27]; while cognitive impairment may predict the presence of TDP-43 burden in the related cognitive region, this is not the case vice versa. Indeed, all language-unimpaired cases in the studied cohort exhibited pTDP-43 pathology to some extent in BA39, often with comparable scoring to language-impaired cases (Table 2).

**Table 2.** Manual TDP43 grading of ALS cases.

| Case | Neuronal (N)<br>Glial (G) | TDP-43 score |  |  |  | ECAS |
| --- | --- | --- | --- | --- | --- | --- |
|  |  | BA4 | BA39 | BA44 | BA46 |  |
| 1 | N | 2 | 1 | 1 | 1 | unimpaired<br>(mismatch) |
|  | G | 2 | 1 | 1 | 1 |  |
| 2 | N | 2 | 1 | 1 | 1 | language |
|  | G | 3 | 1 | 1 | 1 |  |
| 3 | N | 2 | 1 | 1 | 2 | N/A |
|  | G | 2 | 1 | 1 | 1 |  |
| 4 | N | 1 | 1 | 1 | 1 | unimpaired<br>(mismatch) |
|  | G | 2 | 1 | 1 | 2 |  |
| 5 | N | 1 | 0 | 0 | 0 | N/A |
|  | G | 2 | 1 | 0 | 0 |  |
| 6 | N | 1 | 1 | 1 | 1 | N/A |
|  | G | 1 | 0 | 0 | 0 |  |
| 7 | N | 2 | 1 | 1 | 1 | executive (language<br>mismatch) |
|  | G | 0 | 0 | 0 | 0 |  |
| 8 | N | 1 | 1 | 0 | 0 | language |
|  | G | 2 | 0 | 0 | 0 |  |
| 9 | N | 3 | 1 | 1 | 1 | language |
|  | G | 3 | 1 | 1 | 1 |  |
| 10 | N | 2 | 3 | 3 | 3 | all (FTD) |
|  | G | 3 | 3 | 3 | 3 |  |

#### Discussion and Future Work

Few studies have previously investigated markers of neuroinflammation and their relationship with pathological proteins in ALS *post-mortem* tissue[10, 17, 50, 52]. This study is the first to conduct immunohistochemical staining analysis at this scale with unbiased, high-throughput quantification at the image, cell and superpixel level, with diverse regional coverage in deeply clinically phenotyped *post-mortem* tissue. Here we outlined a potential standard for the quantitative assessment of important neuropathological elements in ALS, building upon previous methods analysing FUS- and splicing factor proline- and glutamine-rich (SFPQ)-related differences at the single-cell level in sporadic ALS spinal cord tissue[28], and that described for Alzheimer’s disease[43, 60]. Additionally, this study allowed for *post-mortem* tissue staining data to be directly compared with both motor and cognitive clinical features. Importantly, our random forest model revealed that, from a standard panel of neuropathological markers, the microglia-related stains Iba1 and CD68 were more predictive of C9-ALS status compared to controls than pTDP-43 or GFAP staining. The relevance of microglia in C9-ALS may pertain to the proposed pathogenic mechanism of *C9orf72* HREs being haploinsufficiency. Microglia have previously been shown to strongly express and require C9orf72 protein for normal function, and may therefore be particularly susceptible to the negative effects of this haploinsufficiency[38, 46, 48]. Indeed, *C9orf72* -/- peripheral myeloid cells have been demonstrated to have hyperactive type I interferon responses, mediated by impaired lysosomal degradation of the stimulator of interferon genes (STING) protein[38], thus suggesting that loss of normal C9orf72 function may lead to a proinflammatory state, as we observed in our data. Similarly, mouse *C9orf72* -/- microglia have also been shown to have higher levels of STING protein and increased expression of interferon response genes as well as other activation response genes[32]. Qualitative pathological assessment has shown signs of increased microglial activation in *C9orf72* HRE *post mortem* tissue[18], and microglial activation in the cervical corticospinal tract has been shown to correlate with disease progression[11]. Our study supports and builds upon these findings, highlighting morphological and spatial features of microglia that may be at play, and suggesting an influence of microglial activation in extra-motor regions on cognitive impairment in C9-ALS-FTSD.

There are a few possible reasons as to why pTDP-43 staining was not found to be predictive of disease status in this study. For instance, the way ROIs were selected may lead images from disease cases to appear free of pTDP- 43. This was most evident when comparing automated and manual grading methods, and quantification of pTDP- 43 in brain regions with less aggregate burden (i.e., extra-motor regions) (Figure S3). In line with this, it is recommended that future analyses of pTDP-43 staining combine manual grading of burden with intensity, morphological and spatial features afforded by digital analysis. Alternatively, as has previously been described regarding cognitive impairment[26], it could be that while affected cases all displayed TDP-43 pathology, some unaffected cases, termed “mismatch cases”, could be resilient to TDP-43 burden. Indeed, while TDP-43 pathology has been associated with more rapid cognitive decline[62], it has also been observed in healthy ageing[24, 42]. While it may also be the case that the presence of pathology may precede clinical manifestations, such as in Alzheimer’s disease[31], and that healthy control or “mismatch” cases could have had the potential to develop symptoms later on, this would still indicate that while TDP-43 aggregation may have a central role in disease pathogenesis, its presence or burden may not serve as well as a classifier of C9-ALS in a given time point as other markers such as microglial activation. It may be that different conformations of TDP-43 aggregates not discerned in this study exist that lead to differential behaviour of microglia, similarly to what has been observed with microglial infiltration of amyloid plaques in Alzheimer’s disease[12, 15]. This could explain our findings that language-impaired, BA39 TDP43+ cases have significantly more CD68+ staining than controls, while language- unimpaired cases, which are also BA39 TDP43+, do not (Table 2; Figure 5a-b). Furthermore, there appears to be a difference in the Iba1+ CD68+ relationship between language-impaired and language-unimpaired cases (Figure 5c). Recent studies have shown the immunogenic capacity of TDP-43 and its ability to activate microglia specifically[64, 65], though further research is needed to determine whether certain aggregate subtypes induce a differential response in microglia.

Finally, our random forest model revealed FUS staining to be an accurate classifier of C9-ALS status, and we also found differences in the nuclear/cytoplasmic FUS ratio in glia between control and C9-ALS cases. FUS is an RNA-binding protein that is known to undergo liquid-liquid phase separation (LLPS) and associate with membraneless organelles, some of which are localised to the nucleus such as paraspeckles and nuclear gems[37]. It is also known to autoregulate and increase its own expression as a result of impaired nuclear import[30]. We did not find significant differences in *FUS* expression between control and disease using RT-qPCR, which suggests possible LLPS or aggregation rather than compensatory overexpression. It has been shown that functional C9orf72 protein is required for removal of aggregation-prone FUS protein[14], as well as that C9orf72- related DPRs have been shown to induce aggregation of TDP-43, which similarly to FUS has an intrinsically disordered C-terminal region[16]. Thus, C9orf72 haploinsufficiency due to mutation or the interaction of FUS with DPRs may result in its impaired degradation; our observations of cytoplasmic condensate and paraspeckle formation in C9-ALS in the absence of increased *FUS* expression further support this hypothesis. Whilst this is the first comprehensive, unbiased identification of FUS dysregulation in C9-ALS *post-mortem* tissue, several caveats should be considered. For example, future studies with a higher sample size and cell-sorting methods may be needed to investigate expression changes further, as any cell-type-specific effects may be masked during measurement of bulk RNA expression. Additionally, the cell segmentation algorithm employed in this study was less robust than the superpixel segmentation algorithm; to avoid frequent detection of nonexistent cells over all images, haematoxylin thresholds for the identification of nuclei were set relatively high, resulting in some cells being excluded from the analysis (Figure 1b). Thus, our cell segmentation findings may not be as representative of all cells in the ROIs compared to our superpixel segmentation findings. As such, the question of how FUS staining is predictive of C9-ALS status and subsequent cytotoxic mechanisms requires further investigation.

There are limitations that should be noted regarding all digital analyses. For example, pre-set staining thresholds in automated digital analysis circumvents the possibility of bias that arises with manual grading, though it is not as accommodating of any natural variation in staining. Additionally, this study utilised manually selected ROIs based on predetermined criteria rather than entire tissue scans, which were designed to equally represent different regions of interest as part of an exploratory analysis (i.e., grey and white matter, vascular-adjacent and non-vascular-adjacent regions), though as a result, the analysis risked being less sensitive to sparser pathology. Thus, it may be worth assessing larger regions, such as entire grey and white matter regions within each tissue section, although this would require more sophisticated cortical and cellular annotation algorithms than are currently available. Additionally, while our cohort included deeply clinically phenotyped *post-mortem* tissue and was sufficiently powered to detect differences in staining with regional resolution, our sample size did not include multiple fluency-impaired cases, nor sufficient cases with executive impairment to conduct meaningful comparisons regarding these types of cognitive impairment. Finally, while the random forest model provides important information about which stains are accurate classifiers of C9-ALS, it is currently unclear whether the predictive features are specific to C9-ALS, or whether they are common to other ALS subtypes or to neurodegeneration in general. To determine this, immunohistochemistry features from other neurodegenerative diseases would need to be added to the random forest model.

These limitations notwithstanding, our study demonstrates the importance of a standardised digital pathological assessment of commonly used immunohistochemical markers in ALS and other conditions, the results of which can be shared and meta-analysed to provide a comprehensive and highly powered model revealing pathological predictors of disease and clinical phenotypes. The identification of these predictors may hold promise for peripheral biomarkers and positron emission tomography (PET) ligand creation in the future and contribute to the essential understanding of disease-related specific pathways for the development of targeted therapeutics. Together, our high-throughput digital pathology and random forest modelling approach enabled detailed mapping and analysis of tens of millions of features derived from common neuropathological stains to characterise neuroinflammation and protein dynamics across a clinically heterogeneous C9-ALS cohort. The generation of large datasets using this method allows for the investigation of numerous neuropathological questions, many of which are outside the scope of this study. For this reason, all the scripts and data featured in this study are included in Supplementary Materials.

## Acknowledgments

This research was funded in part by a studentship from the Wellcome Trust (108890/Z/15/Z) to OMR and MDES, a Pathological Society and Jean Shanks foundation grant (217CHA R46564) to JMG and JO, and a Sir Henry Dale fellowship jointly funded by the Wellcome Trust and the Royal Society (215454/Z/19/Z) to CRS. We gratefully acknowledge Dr. Tom Gillingwater for his helpful comments and support. This work would also not be possible without the resources of the Edinburgh Brain Bank. The authors declare no conflicts of interest. SD numbers of cases from the Edinburgh Brain Bank included in the study are available upon request.

## Notes

https://figshare.com/articles/figure/C9-ALS_and_control_images/17145896

https://doi.org/10.6084/m9.figshare.17145902.v1

https://figshare.com/articles/dataset/Random_forest_results/17145905

### Competing Interest Statement

The authors have declared no competing interest.

