## Supplementary Materials for "Random forest modelling of neuropathological features identifies microglial activation as an accurate pathological classifier of C9orf72-related amyotrophic lateral sclerosis"

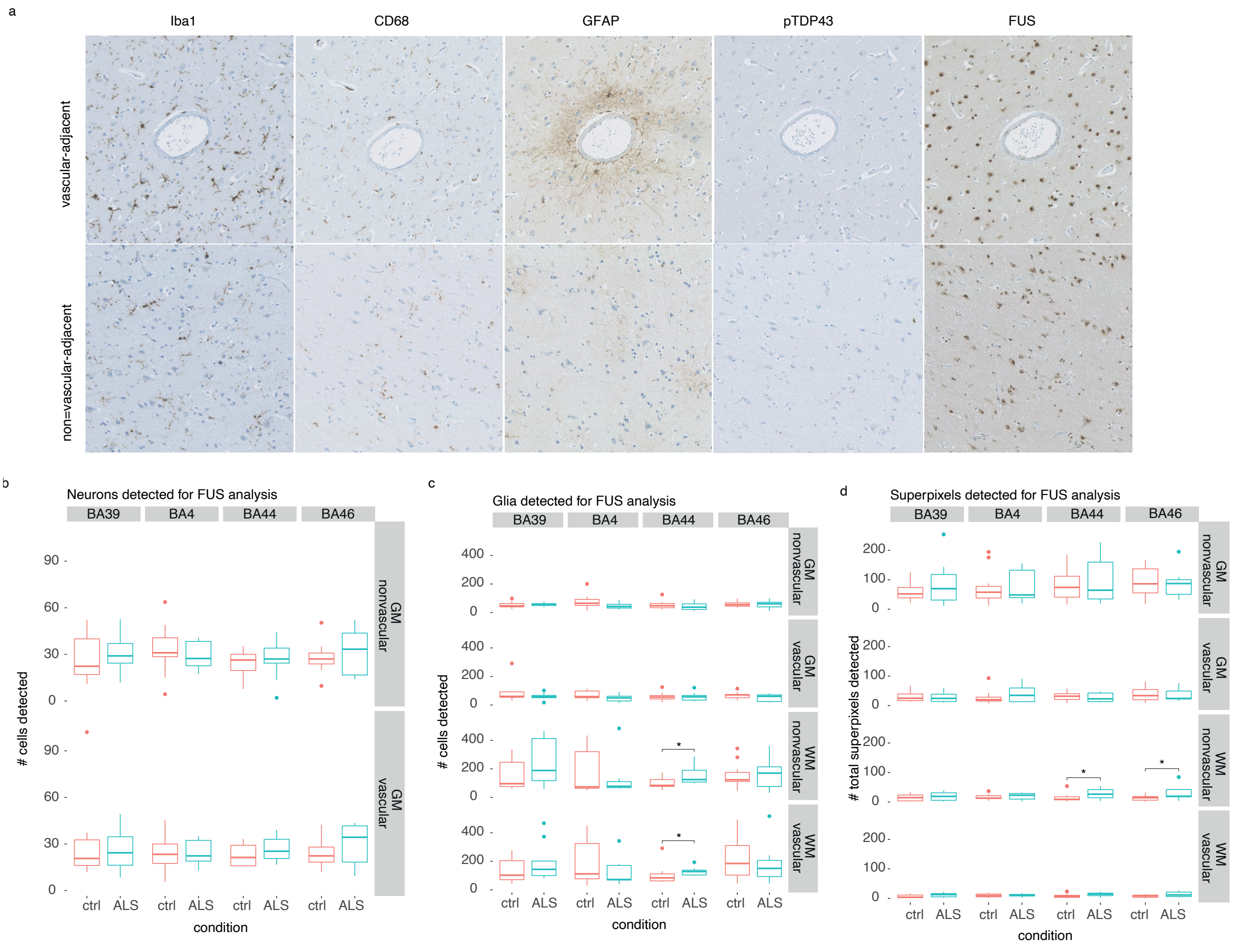


**Supplementary Figure 1.** ***Supplementary digital pathology analysis information.*** (a) Immunohistochemical stains included in digital pathology analysis, showing (top) vascular-adjacent and (bottom) non-vascular adjacent regions. (b) Box and whisker plot showing the number of neurons detected in FUS cell segmentation analysis across all included regions. (c) Box and whisker plot showing the number of glia detected in FUS cell segmentation analysis across all included regions. (d) Box and whisker plot showing the total number of superpixels detected in FUS superpixel segmentation analysis across all included regions. Data in box and whisker plots are averaged across quantified regions not included in the presented stratification, such that each category contains n = number of cases, to avoid pseudoreplication. BA, Brodmann area; GM, grey matter; WM, white matter. Mann-Whitney U test, * $p < 0.05$, ** $p < 0.01$, *** $p < 0.001$, **** $p < 0.0001$.

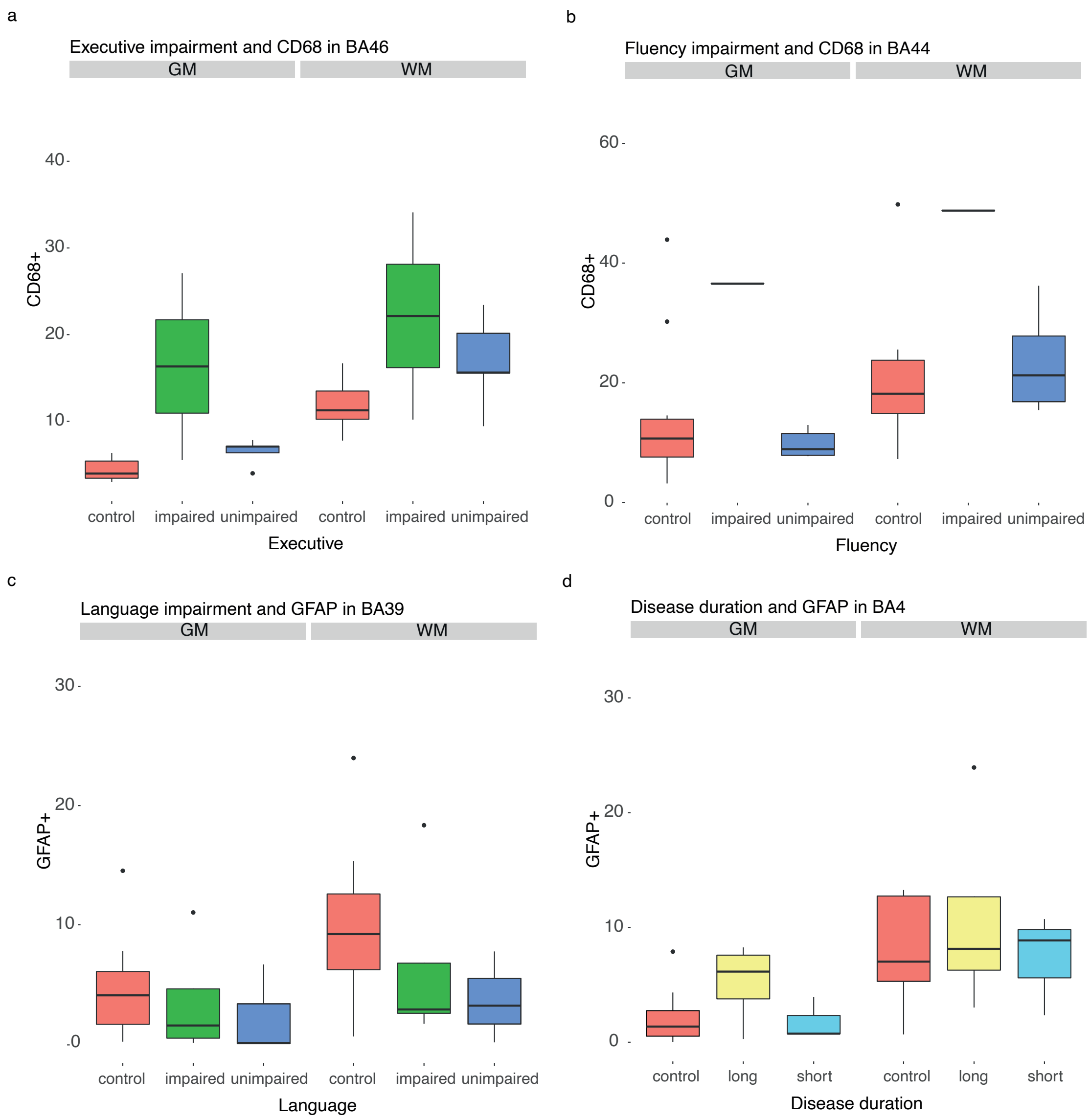


**Supplementary Figure 2.** ***Additional glial activation staining in cognitive regions regions.*** (a) Box and whisker plot of CD68+ superpixel staining in BA46 between control, executive-impaired and executive-unimpaired cases. (e) Box and whisker plot of CD68+ superpixel staining in BA44 between control, fluency-unimpaired case and one fluency-impaired (FTD) case. No statistical tests were conducted for (a) & (b) due to low sample size. (c) Box and whisker plot of GFAP+ superpixel staining in BA39 between control, language-impaired and language-unimpaired cases. (d) Box and whisker plot of GFAP+ superpixel staining in BA4 between control, long and short survivors. Wilcoxon tests with Holm-Sidak multiple comparisons correction were conducted for (c) & (d), with no significant differences found. Data in box and whisker plots are averaged across quantified regions not included in the presented stratification, such that each category contains n = number of cases, in order to avoid pseudoreplication. BA, Brodmann area; GM, grey matter; WM, white matter.

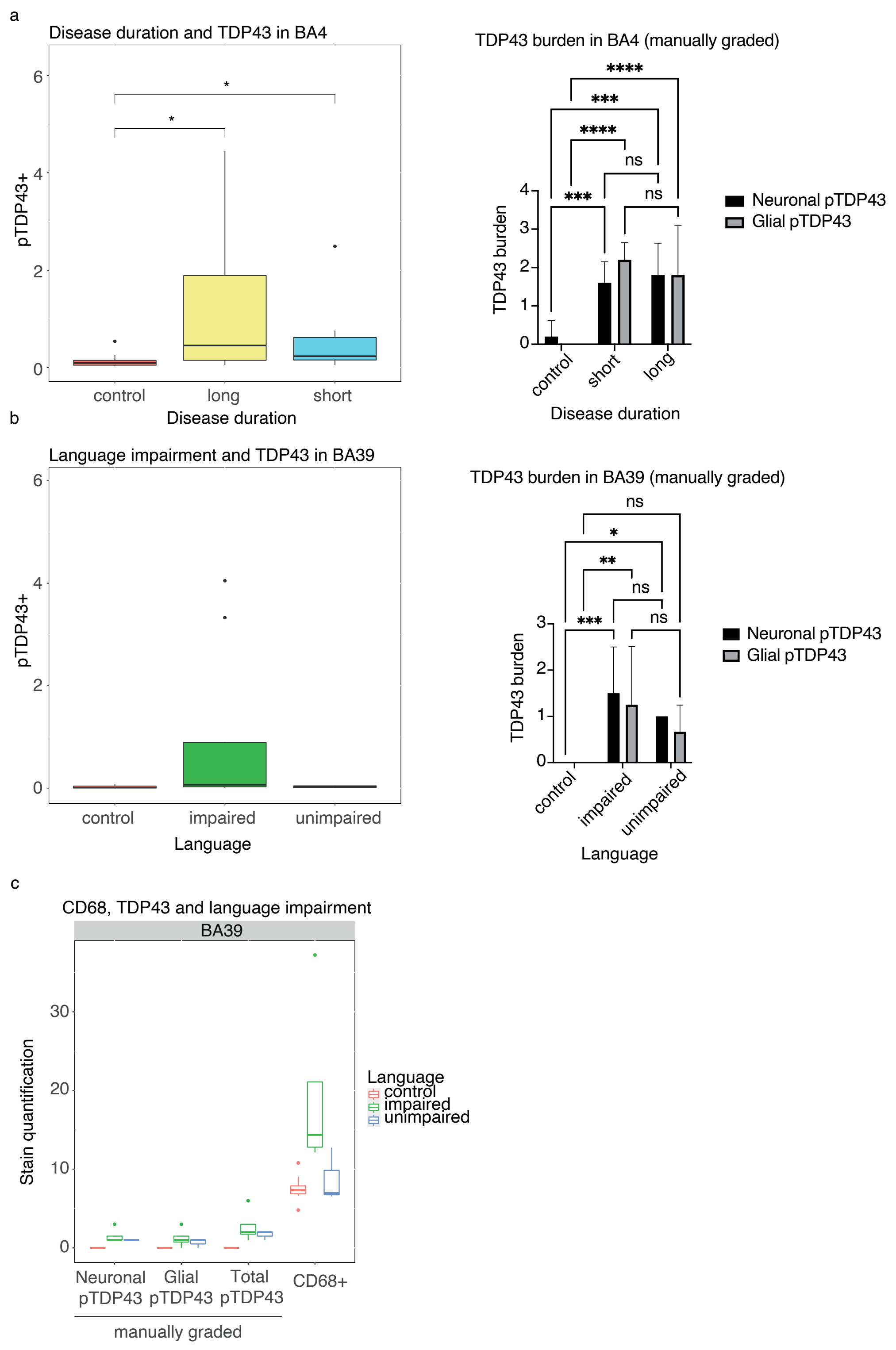


**Supplementary Figure 3.** ***Comparison of automated digital pathology analysis and manual grading methods of TDP-43 burden quantification.*** (a & b) Box and whisker plots showing the comparison between digital analysis (Wilcoxon, Holm-Sidak test) (left) and manual pTDP-43 grading (ANOVA, Tukey's test) by a blinded pathologist (right) in BA39 in cases stratified by language impairment. (c) Box and whisker plot showing size of differences in manual pTDP-43 grading and digitally quantified CD68+ superpixels between language-impaired and language-unimpaired ALS cases. Data in box and whisker plots are averaged across quantified regions not included in the presented stratification, such that each category contains n = number of cases to avoid pseudoreplication. BA, Brodmann area. * $p < 0.05$, ** $p < 0.01$, *** $p < 0.001$, **** $p < 0.0001$.

| by | condition | TRUE | FALSE | EXPECTED | p-value | sensitivity | specificity |
|---|---|---|---|---|---|---|---|
| all | control | 527 | 273 | 400 | p < 0.0001 | 0.69 | 0.66 |
| | disease | 551 | 249 | 400 | p < 0.0001 | | |
| CD68 | control | 96 | 64 | 80 | p = 0.0114 | 0.67 | 0.60 |
| | disease | 107 | 53 | 80 | p < 0.0001 | | |
| FUS | control | 111 | 49 | 80 | p < 0.0001 | 0.64 | 0.69 |
| | disease | 103 | 57 | 80 | p = 0.0003 | | |
| GFAP | control | 84 | 76 | 80 | p = 0.527 | 0.79 | 0.53 |
| | disease | 127 | 33 | 80 | p < 0.0001 | | |
| Iba1 | control | 134 | 26 | 80 | p < 0.0001 | 0.78 | 0.84 |
| | disease | 125 | 35 | 80 | p < 0.0001 | | |
| pTDP43 | control | 102 | 58 | 80 | p = 0.0005 | 0.51 | 0.64 |
| | disease | 81 | 79 | 80 | p = 0.8744 | | |
| BA4 | control | 122 | 78 | 100 | p = 0.0019 | 0.79 | 0.61 |
| | disease | 158 | 42 | 100 | p < 0.0001 | | |
| BA39 | control | 136 | 64 | 100 | p < 0.0001 | 0.64 | 0.68 |
| | disease | 127 | 73 | 100 | p = 0.0001 | | |
| BA44 | control | 136 | 64 | 100 | p < 0.0001 | 0.63 | 0.68 |
| | disease | 126 | 74 | 100 | p = 0.0002 | | |
| BA46 | control | 133 | 67 | 100 | p < 0.0001 | 0.66 | 0.67 |
| | disease | 132 | 68 | 100 | p < 0.0001 | | |
| GM | control | 271 | 129 | 200 | p < 0.0001 | 0.68 | 0.68 |
| | disease | 273 | 127 | 200 | p < 0.0001 | | |
| WM | control | 256 | 144 | 200 | p < 0.0001 | 0.68 | 0.64 |
| | disease | 270 | 130 | 200 | p < 0.0001 | | |
| VA | control | 268 | 132 | 200 | p < 0.0001 | 0.65 | 0.67 |
| | disease | 260 | 140 | 200 | p < 0.0001 | | |
| NVA | control | 259 | 141 | 200 | p < 0.0001 | 0.71 | 0.65 |
| | disease | 283 | 117 | 200 | p < 0.0001 | | |

**Supplementary Table 1. Random forest classification of stain panel with sensitivity and specificity scores.** Results were considered significant if the number classified correctly was significantly better than chance according to $X^2$ tests.

| condition | TRUE | FALSE | Iteration | feature | %improvement |
|---|---|---|---|---|---|
| disease | 542 | 258 | 36 | 2++Mean(Min.diameter.px)_detections | -0.1841621 |
| disease | 542 | 258 | 45 | 3++Mean(Max.diameter.px)_detections | -0.1841621 |
| disease | 542 | 258 | 59 | Negative+Mean(Distance.to.detection.with.2+.px)_detections | -0.1841621 |
| disease | 542 | 258 | 88 | glia+Mean(Nucleus..Hematoxylin.OD.range) | -0.1841621 |
| disease | 542 | 258 | 89 | glia+Mean(Nucleus..DAB.OD.mean) | -0.1841621 |
| disease | 542 | 258 | 90 | glia+Mean(Nucleus..DAB.OD.sum) | -0.1841621 |
| disease | 542 | 258 | 92 | glia+Mean(Nucleus..DAB.OD.max) | -0.1841621 |
| disease | 542 | 258 | 95 | glia+Mean(Cell..Area) | -0.1841621 |
| disease | 542 | 258 | 147 | neuron+Mean(Cell..DAB.OD.mean) | -0.1841621 |
| disease | 541 | 259 | 37 | 2++Mean(Distance.to.detection.with.Negative.px)_detections | -0.3683241 |
| disease | 541 | 259 | 43 | 3++Mean(Circularity)_detections | -0.3683241 |
| disease | 541 | 259 | 85 | glia+Mean(Nucleus..Hematoxylin.OD.std.dev) | -0.3683241 |
| disease | 541 | 259 | 116 | glia+Mean(Cytoplasm..DAB.OD.min) | -0.3683241 |
| disease | 541 | 259 | 117 | glia+Mean(Nucleus.Cell.area.ratio) | -0.3683241 |
| disease | 541 | 259 | 140 | neuron+Mean(Cell..Max.caliper) | -0.3683241 |
| disease | 541 | 259 | 142 | neuron+Mean(Cell..Eccentricity) | -0.3683241 |
| disease | 541 | 259 | 153 | neuron+Mean(Cytoplasm..Hematoxylin.OD.max) | -0.3683241 |
| disease | 541 | 259 | 160 | neuron+Mean(nuc_cyt_intensity_ratio) | -0.3683241 |
| disease | 540 | 260 | 4 | Allred score_polygon | -0.5524862 |
| disease | 540 | 260 | 40 | 2++Mean(Distance.to.detection.with.3+.px)_detections | -0.5524862 |
| disease | 540 | 260 | 63 | 3++Mean(norm_dist_vessel)_detections | -0.5524862 |
| disease | 540 | 260 | 71 | Retract Area_retraction | -0.5524862 |
| disease | 540 | 260 | 91 | glia+Mean(Nucleus..DAB.OD.std.dev) | -0.5524862 |
| disease | 539 | 261 | 18 | %2+_annotation | -0.7366483 |
| disease | 539 | 261 | 23 | 1++Mean(Circularity)_detections | -0.7366483 |
| disease | 539 | 261 | 54 | Negative+Mean(Solidity)_detections | -0.7366483 |
| disease | 539 | 261 | 61 | 1++Mean(norm_dist_vessel)_detections | -0.7366483 |
| disease | 539 | 261 | 110 | glia+Mean(Cytoplasm..Hematoxylin.OD.std.dev) | -0.7366483 |
| disease | 539 | 261 | 113 | glia+Mean(Cytoplasm..DAB.OD.mean) | -0.7366483 |
| disease | 539 | 261 | 115 | glia+Mean(Cytoplasm..DAB.OD.max) | -0.7366483 |
| disease | 539 | 261 | 141 | neuron+Mean(Cell..Min.caliper) | -0.7366483 |
| disease | 539 | 261 | 152 | neuron+Mean(Cytoplasm..Hematoxylin.OD.std.dev) | -0.7366483 |
| disease | 538 | 262 | 12 | Allred proportion_annotation | -0.9208103 |
| disease | 538 | 262 | 30 | 1++Mean(Distance.to.detection.with.3+.px)_detections | -0.9208103 |
| disease | 538 | 262 | 42 | 3++Mean(Length.px)_detections | -0.9208103 |
| disease | 538 | 262 | 84 | glia+Mean(Nucleus..Hematoxylin.OD.sum) | -0.9208103 |
| disease | 538 | 262 | 93 | glia+Mean(Nucleus..DAB.OD.min) | -0.9208103 |
| disease | 538 | 262 | 98 | glia+Mean(Cell..Max.caliper) | -0.9208103 |
| disease | 538 | 262 | 101 | glia+Mean(Cell..Hematoxylin.OD.mean) | -0.9208103 |
| disease | 538 | 262 | 107 | glia+Mean(Cell..DAB.OD.max) | -0.9208103 |
| disease | 538 | 262 | 108 | glia+Mean(Cell..DAB.OD.min) | -0.9208103 |
| disease | 538 | 262 | 112 | glia+Mean(Cytoplasm..Hematoxylin.OD.min) | -0.9208103 |
| disease | 538 | 262 | 132 | neuron+Mean(Nucleus..DAB.OD.sum) | -0.9208103 |
| disease | 538 | 262 | 134 | neuron+Mean(Nucleus..DAB.OD.max) | -0.9208103 |
| disease | 537 | 263 | 7 | %1+_polygon | -1.1049724 |
| disease | 537 | 263 | 24 | 1++Mean(Solidity)_detections | -1.1049724 |
| disease | 537 | 263 | 31 | 2++Mean(Area.px^2)_detections | -1.1049724 |
| disease | 537 | 263 | 35 | 2++Mean(Max.diameter.px)_detections | -1.1049724 |
| disease | 537 | 263 | 49 | 3++Mean(Distance.to.detection.with.2+.px)_detections | -1.1049724 |
| disease | 537 | 263 | 50 | 3++Mean(Distance.to.detection.with.3+.px)_detections | -1.1049724 |
| disease | 537 | 263 | 53 | Negative+Mean(Circularity)_detections | -1.1049724 |
| disease | 537 | 263 | 58 | Negative+Mean(Distance.to.detection.with.1+.px)_detections | -1.1049724 |
| disease | 537 | 263 | 66 | H-score_retraction | -1.1049724 |
| disease | 537 | 263 | 83 | glia+Mean(Nucleus..Hematoxylin.OD.mean) | -1.1049724 |
| disease | 537 | 263 | 86 | glia+Mean(Nucleus..Hematoxylin.OD.max) | -1.1049724 |

| | | | | | |
|---|---|---|---|---|---|
| disease | 537 | 263 | 106 | glia+Mean(Cell..DAB.OD.std.dev) | -1.1049724 |
| disease | 537 | 263 | 127 | neuron+Mean(Nucleus..Hematoxylin.OD.std.dev) | -1.1049724 |
| disease | 537 | 263 | 148 | neuron+Mean(Cell..DAB.OD.std.dev) | -1.1049724 |
| disease | 537 | 263 | 157 | neuron+Mean(Cytoplasm..DAB.OD.max) | -1.1049724 |
| disease | 536 | 264 | 1 | H-score_polygon | -1.2891344 |
| disease | 536 | 264 | 8 | %2+_polygon | -1.2891344 |
| disease | 536 | 264 | 11 | H-score_annotation | -1.2891344 |
| disease | 536 | 264 | 19 | %3+_annotation | -1.2891344 |
| disease | 536 | 264 | 25 | 1++Mean(Max.diameter.px)_detections | -1.2891344 |
| disease | 536 | 264 | 47 | 3++Mean(Distance.to.detection.with.Negative.px)_detections | -1.2891344 |
| disease | 536 | 264 | 48 | 3++Mean(Distance.to.detection.with.1+.px)_detections | -1.2891344 |
| disease | 536 | 264 | 68 | Allred intensity_retraction | -1.2891344 |
| disease | 536 | 264 | 70 | Vessel Area_retraction | -1.2891344 |
| disease | 536 | 264 | 72 | Retraction %_retraction | -1.2891344 |
| disease | 536 | 264 | 100 | glia+Mean(Cell..Eccentricity) | -1.2891344 |
| disease | 536 | 264 | 120 | neuron+Mean(Nucleus..Perimeter) | -1.2891344 |
| disease | 536 | 264 | 121 | neuron+Mean(Nucleus..Circularity) | -1.2891344 |
| disease | 536 | 264 | 129 | neuron+Mean(Nucleus..Hematoxylin.OD.min) | -1.2891344 |
| disease | 536 | 264 | 154 | neuron+Mean(Cytoplasm..Hematoxylin.OD.min) | -1.2891344 |
| disease | 535 | 265 | 0 | Positive %_polygon | -1.4732965 |
| disease | 535 | 265 | 3 | Allred intensity_polygon | -1.4732965 |
| disease | 535 | 265 | 22 | 1++Mean(Length.px)_detections | -1.4732965 |
| disease | 535 | 265 | 41 | 3++Mean(Area.px^2)_detections | -1.4732965 |
| disease | 535 | 265 | 51 | Negative+Mean(Area.px^2)_detections | -1.4732965 |
| disease | 535 | 265 | 55 | Negative+Mean(Max.diameter.px)_detections | -1.4732965 |
| disease | 535 | 265 | 73 | %1+_retraction | -1.4732965 |
| disease | 535 | 265 | 77 | glia+Mean(Nucleus..Area) | -1.4732965 |
| disease | 535 | 265 | 81 | glia+Mean(Nucleus..Min.caliper) | -1.4732965 |
| disease | 535 | 265 | 82 | glia+Mean(Nucleus..Eccentricity) | -1.4732965 |
| disease | 535 | 265 | 97 | glia+Mean(Cell..Circularity) | -1.4732965 |
| disease | 535 | 265 | 126 | neuron+Mean(Nucleus..Hematoxylin.OD.sum) | -1.4732965 |
| disease | 535 | 265 | 128 | neuron+Mean(Nucleus..Hematoxylin.OD.max) | -1.4732965 |
| disease | 535 | 265 | 130 | neuron+Mean(Nucleus..Hematoxylin.OD.range) | -1.4732965 |
| disease | 535 | 265 | 131 | neuron+Mean(Nucleus..DAB.OD.mean) | -1.4732965 |
| disease | 534 | 266 | 6 | Perimeter px_polygon | -1.6574586 |
| disease | 534 | 266 | 9 | %3+_polygon | -1.6574586 |
| disease | 534 | 266 | 15 | Area px^2_annotation | -1.6574586 |
| disease | 534 | 266 | 16 | Perimeter px_annotation | -1.6574586 |
| disease | 534 | 266 | 26 | 1++Mean(Min.diameter.px)_detections | -1.6574586 |
| disease | 534 | 266 | 57 | Negative+Mean(Distance.to.detection.with.Negative.px)_detections | -1.6574586 |
| disease | 534 | 266 | 75 | %3+_retraction | -1.6574586 |
| disease | 534 | 266 | 79 | glia+Mean(Nucleus..Circularity) | -1.6574586 |
| disease | 534 | 266 | 102 | glia+Mean(Cell..Hematoxylin.OD.std.dev) | -1.6574586 |
| disease | 534 | 266 | 104 | glia+Mean(Cell..Hematoxylin.OD.min) | -1.6574586 |
| disease | 534 | 266 | 114 | glia+Mean(Cytoplasm..DAB.OD.std.dev) | -1.6574586 |
| disease | 534 | 266 | 135 | neuron+Mean(Nucleus..DAB.OD.min) | -1.6574586 |
| disease | 534 | 266 | 137 | neuron+Mean(Cell..Area) | -1.6574586 |
| disease | 534 | 266 | 149 | neuron+Mean(Cell..DAB.OD.max) | -1.6574586 |
| disease | 534 | 266 | 158 | neuron+Mean(Cytoplasm..DAB.OD.min) | -1.6574586 |
| disease | 533 | 267 | 2 | Allred proportion_polygon | -1.8416206 |
| disease | 533 | 267 | 29 | 1++Mean(Distance.to.detection.with.2+.px)_detections | -1.8416206 |
| disease | 533 | 267 | 38 | 2++Mean(Distance.to.detection.with.1+.px)_detections | -1.8416206 |
| disease | 533 | 267 | 52 | Negative+Mean(Length.px)_detections | -1.8416206 |
| disease | 533 | 267 | 151 | neuron+Mean(Cytoplasm..Hematoxylin.OD.mean) | -1.8416206 |
| disease | 533 | 267 | 156 | neuron+Mean(Cytoplasm..DAB.OD.std.dev) | -1.8416206 |
| disease | 532 | 268 | 5 | Area px^2_polygon | -2.0257827 |
| disease | 532 | 268 | 27 | 1++Mean(Distance.to.detection.with.Negative.px)_detections | -2.0257827 |
| disease | 532 | 268 | 28 | 1++Mean(Distance.to.detection.with.1+.px)_detections | -2.0257827 |
| disease | 532 | 268 | 56 | Negative+Mean(Min.diameter.px)_detections | -2.0257827 |

| | | | | | |
|---|---|---|---|---|---|
| disease | 532 | 268 | 74 | %2+_retraction | -2.0257827 |
| disease | 532 | 268 | 78 | glia+Mean(Nucleus..Perimeter) | -2.0257827 |
| disease | 532 | 268 | 80 | glia+Mean(Nucleus..Max.caliper) | -2.0257827 |
| disease | 532 | 268 | 99 | glia+Mean(Cell..Min.caliper) | -2.0257827 |
| disease | 532 | 268 | 122 | neuron+Mean(Nucleus..Max.caliper) | -2.0257827 |
| disease | 531 | 269 | 10 | Positive %_annotation | -2.2099448 |
| disease | 531 | 269 | 13 | Allred intensity_annotation | -2.2099448 |
| disease | 531 | 269 | 65 | Positive %_retraction | -2.2099448 |
| disease | 531 | 269 | 109 | glia+Mean(Cytoplasm..Hematoxylin.OD.mean) | -2.2099448 |
| disease | 531 | 269 | 119 | neuron+Mean(Nucleus..Area) | -2.2099448 |
| disease | 530 | 270 | 14 | Allred score_annotation | -2.3941068 |
| disease | 530 | 270 | 33 | 2++Mean(Circularity)_detections | -2.3941068 |
| disease | 530 | 270 | 105 | glia+Mean(Cell..DAB.OD.mean) | -2.3941068 |
| disease | 530 | 270 | 118 | glia+Mean(nuc_cyt_intensity_ratio) | -2.3941068 |
| disease | 529 | 271 | 123 | neuron+Mean(Nucleus..Min.caliper) | -2.5782689 |
| disease | 529 | 271 | 136 | neuron+Mean(Nucleus..DAB.OD.range) | -2.5782689 |
| disease | 528 | 272 | 34 | 2++Mean(Solidity)_detections | -2.7624309 |
| disease | 528 | 272 | 69 | Allred score_retraction | -2.7624309 |
| disease | 528 | 272 | 103 | glia+Mean(Cell..Hematoxylin.OD.max) | -2.7624309 |
| disease | 527 | 273 | 17 | %1+_annotation | -2.946593 |
| disease | 527 | 273 | 67 | Allred proportion_retraction | -2.946593 |
| disease | 525 | 275 | 64 | Negative+Mean(norm_dist_vessel)_detections | -3.3149171 |
| disease | 525 | 275 | 124 | neuron+Mean(Nucleus..Eccentricity) | -3.3149171 |
| disease | 524 | 276 | 125 | neuron+Mean(Nucleus..Hematoxylin.OD.mean) | -3.4990792 |

**Supplementary Table 2. Random forest leave-one-out feature analysis.** List of features that when left out resulted in worse predictivity of disease status.

**Supplementary Information**

Link to raw images
https://figshare.com/articles/figure/C9-ALS_and_control_images/17145896

Link to digital pathology features
https://doi.org/10.6084/m9.figshare.17145902.v1

Link to random forest results
https://figshare.com/articles/dataset/Random_forest_results/17145905

### Supplementary Methods

*FUS RT-qPCR*

Primers that failed to amplify FUS in FFPE tissue samples, targeting two non-contiguous central regions (R1, R2) listed below. Note that R2P3 showed some amplification but was not as consistent between triplicates as the primer pair used in the study (R2P2).

Region 1

```
GTTCACACGTGGTGGCATGAAAAGAGTGGCTAAAGTGGTATCAAGACTGCCTGGATGTTCTTTGAAACTA
TTATAAAAAGGAAACTGAAAAAAATGGGGATAGAGAAGGAAGGGAGTTAGGTGTGTCCTTAGTTAGCAGT
GAGAAGTATTTGTTACGAAGTATTTCTCAGAAATACCTGGCTTGTGGGTTCCACCCCCAGTGATTTAGGT
CTGAGAGGACCCTGAAAATCTACCTTTCTAACAAGTCCCCAGTGATGCTGATGCGTCTGGACCACACTCA
GATGGTTTACAGCAGTGGTTCTTTCAAAATGTGGATCATGTCCAAGTTGGTGAACAAAACTAGTGAAAGA
CCTGACCACATGAAGTAAAGCATTGAACTCCTGTTTAGGTTGTATTGATGTTTGTGTACTAGATTTGAAT
GTAAAATGGGTTTCTTATTTAATTCGGGGACCTTCAACTGAAAGTTGATAAATACTGATGGACTTTTCTG
TGTGGTCTTGTTGGGCATTTCACTCCTGAGCCCTGGTTTCCATACTGTATACTGGGTGTTAACATGTTTC
AAAGGATAATTGTCAAACTGAATCTGAAATTTATCAGCATGGCTGGCATATAGGGACTCAAAAGGGATGT
GGATTTCTTTTTAGTTGTCTTCCATAAACCAAATGATACCAGTTGCTTGATGGATACTAGGTGCTTTAGG
TTTTTTCCTGTGTTTTTTATTTTACCTTTTCACATTTGCATTTTCTCTGTTCAACAAGCAGAACAGGATA
ATTCAGACAACAACACCATCTTTGTGCAAGGCCTGGGTGAGAATGTTACAATTGAGTCTGTGGCTGATTA
CTTCAAGCAGATTGGTATTATTAAGGTACTTGTGGAGAGGAGTGGGAGCTTTCTGTCAGTGTTGTAGGCT
TGTGGATTTCACACATTAGTAAAAGCAAGTCTTTAATGGTTGCCAGCAGTAAAAACAAGTCTTAGTGGTT
GTTGCCAGCTTAATTTGTTGAGGAAAGAGCCTTAGTTACTGTTTTCTAAAAGAGAAGTTCTATCTTAACA
CAAAAAGTATAACTTATCAGAGTACCCTAAACTCTTGAGATTTGTACTCTATAGTAACTTTTAGTTTTAT
CTTTCAATATTGGAGTGAGAGACAGTTTTCTTTAATGGAGGTTTACATGTGAGGTAGGAAGAAGTAACTG
GGAAGAGGGGAGCTGAAGTTTGGGAATTATAAACCTCATGTTCTAGAGGAAGAAGATGGAAAGGGAGTAC
TGTAGCCTTTAAAATTGATGTTACCTCATTTTGCTTTCTTCAGACAAACAAGAAAACGGGACAGCCCATG
ATTAATTTGTACACAGACAGGGAAACTGGCAAGCTGAAGGGAGAGGCAACGGTCTCTTTTGATGACCCAC
CTTCAGCTAAAGCAGCTATTGACTGGTTTGATGGTATGTATGAGAAGGCTGGCAGAGGTGGGGCTGGGGA
TATAGGGCAGCAAGCCTTAGGAAACAAGCCATAGTTTGAGGGTTCTTTTGAGTCTTCCAACACTTACTTT
```

Region 2

```
GTTGGCCAGGCTGGTCTCGAACTCCTGACCTCGTGATCCACACGCCTCGGCCTCCCAAAGTGCTGGTATT
ACAGGCGTGAGCCACCGCACCCGGTCAGCTTTATTATTAAATGTTAACTTCATCTGCTTTGTACACGAAT
GCATACCCAGTGCCCAGAACAGTGCCTGGAACATACTAGGTGCTAAATAAATATTTGTGACTTAATGCAT
GAATAAGGGTGGACTTCCTTTCTTTTGCTCTCACTGGAGAGTTGAACTCTCCTTCCAAAGGCGGTGGGGT
GGATATTGGCATATTCAGGGCCTTTAGGGCTGAAGTCAAGGGCTTAGTGGGGCTTAATTTGTGGGCGGGC
CCAGGGCATGGCCCTCATTGTTTCTCTAGAAAGACGTGTCCAACCCTCAAAGGACCTTCTGAAATCCCGC
TGAAAGGTTAAGTTGGGAAGGAAAACCTGCATGCCTATGTATCAGGAATTAACGTCTTTTGTCTTGTTTT
ATTCAGTAGTTTCAAATTGCCTTCTCCAGGCAAGGCTGATGAAAGTGCAAGTTGCAAGTTAATTTGAATG
TTTCTTTTTGCTTTTGCTCTCACAGGAAGTGAAAAGGCGACCAAACACTCTAGCATTCATGCCACCAAAA
AGAGGAGTGTTTTGCAGTTACAAGACCTGGATTCGAATCACGACTCCTCTTAGCTGCCCTGTAATCAGGC
ACAATTACTTGGGTCTCTGAGTCTCACTTTCCTTATCTAGAAAACGGAGGTATCTTTACTTCCTTCGTAA
GACTGATGACAAGGAAATTATCTGTGCATTTTGAAACCACTTAAGCCTTGTACACGTTTTATTTCTGGGA
TCGCCCTGGTAGGGCTTCAGAAAAATAAAAAGGAGGTCCCTGAGAAAAGGCTGGGTACCGTACATCTGAG
GTCAACCCTCTCTGGTCCCAAGGATGGCCTGGGCTGTTCCGCCCCGTGGCTCCCCAGGGGCAAAGCCATG
AGGATCCGGGTGAGAGCCCAGTGCTGGACGAGCCCGGGGCCCAGGGGTCCCGGCCGAAATCCCTGCTGTC
TTTCAGGTCAAACGTCATAATCCCCGAACCCCAGAAAGGCCGAAAGGCAAGGCAACCCTGAAAGACGACG
AAGTCAACCTCAGGGCGCAGGAGAGGGAGGGCCAGTGTGCTGCCGACGAGGGAGGCTGGAGCCGCGGGGA
CGAGGCGCCCCATACAGCGGCAAGAGGGTGGAGGGCAGGAGCTCGCCATCCTGGGTGAAAGCGGGGCCCA
GCGAAGGGGCCCGGCCACAGGAATCTCGGTTCCACCCCGCTACTCCCGGCTGTGACTCCAGTTTCGTCCC
CAGCCGCCGGGACCGCCCCCTCGCCCCGCCCCCAGCGGGCACTCAGGCCGTACCACTGTGCCTTCATGGG
```

R1P1 Forward: 5’-TGGGGATAGAGAAGGAAGGGAG-3’
R1P1 Reverse: 5’-TCCAGACGCATCAGCATCAC-3’

R1P2 Forward: 5’-AATGGGGATAGAGAAGGAAGG-3’
R1P2 Reverse: 5’-AAGAACCACTGCTGTAAACC-3’

R1P3 Forward: 5’-TGGGGATAGAGAAGGAAGGG-3’
R1P3 Reverse: 5’-AAGAACCACTGCTGTAAACC-3’

R2P1 Forward: 5’-ACTTCCTTTCTTTTGCTCTCAC-3’
R2P1 Reverse: 5’-TTCCTTCCCAACTTAACCTTTC-3’

R2P3 Forward: 5’-ACTTCCTTTCTTTTGCTCTCAC-3’
R2P3 Reverse: 5’-TAAGCCCTTGACTTCAGCC-3’

*Scripts used to generate data*

CD68 NVA

```
setImageType('BRIGHTFIELD_H_DAB');
setColorDeconvolutionStains('{"Name" : "H-DAB default", "Stain 1" : "Hematoxylin", "Values 1" :
"0.65111 0.70119 0.29049 ", "Stain 2" : "DAB", "Values 2" : "0.26917 0.56824 0.77759 ",
"Background" : " 255 255 255 "}');
createSelectAllObject(true);
selectAnnotations ();
runPlugin('qupath.imagej.superpixels.DoGSuperpixelsPlugin', '{"downsampleFactor": 1.0,
"sigmaPixels": 5.0,  "minThreshold": 10.0,  "maxThreshold": 230.0,  "noiseThreshold": 1.0}');
selectDetections();
runPlugin('qupath.lib.algorithms.IntensityFeaturesPlugin', '{"downsample": 1.0,  "region": "ROI",
"tileSizePixels": 200.0,  "colorOD": false,  "colorStain1": false,  "colorStain2": true,  "colorStain3": false,
"colorRed": false,  "colorGreen": false,  "colorBlue": false,  "colorHue": false,  "colorSaturation": false,
"colorBrightness": false,  "doMean": true,  "doStdDev": true,  "doMinMax": true,  "doMedian": true,
"doHaralick": false,  "haralickDistance": 1,  "haralickBins": 32}');
addShapeMeasurements("AREA", "LENGTH", "CIRCULARITY", "SOLIDITY", "MAX_DIAMETER",
"MIN_DIAMETER")
setDetectionIntensityClassifications("ROI: 1.00 px per pixel: DAB: Mean", 0.0348, 0.0693, 0.1037)
detectionCentroidDistances(true)
```

CD68 VA

```
setImageType('BRIGHTFIELD_H_DAB');
setColorDeconvolutionStains('{"Name" : "H-DAB default", "Stain 1" : "Hematoxylin", "Values 1" :
"0.65111 0.70119 0.29049 ", "Stain 2" : "DAB", "Values 2" : "0.26917 0.56824 0.77759 ",
"Background" : " 255 255 255 "}');
selectAnnotations ();
def region = getPathClass('Other')
def other = getPathClass('Region*')
getAnnotationObjects().eachWithIndex { annotation , i ->
   if (i % 2 == 0)
      annotation.setPathClass(region)
  else
      annotation.setPathClass(other)
}
fireHierarchyUpdate()
makeInverseAnnotation()
runPlugin('qupath.imagej.superpixels.DoGSuperpixelsPlugin', '{"downsampleFactor": 1.0,
"sigmaPixels": 5.0,  "minThreshold": 10.0,  "maxThreshold": 230.0,  "noiseThreshold": 1.0}');
selectDetections()
```

```
runPlugin('qupath.lib.algorithms.IntensityFeaturesPlugin', '{"downsample": 1.0,  "region": "ROI",
"tileSizePixels": 200.0,  "colorOD": false,  "colorStain1": false,  "colorStain2": true,  "colorStain3": false,
"colorRed": false,  "colorGreen": false,  "colorBlue": false,  "colorHue": false,  "colorSaturation": false,
"colorBrightness": false,  "doMean": true,  "doStdDev": true,  "doMinMax": true,  "doMedian": true,
"doHaralick": false,  "haralickDistance": 1,  "haralickBins": 32}');
addShapeMeasurements("AREA", "LENGTH", "CIRCULARITY", "SOLIDITY", "MAX_DIAMETER",
"MIN_DIAMETER")
setDetectionIntensityClassifications("ROI: 1.00 px per pixel: DAB: Mean", 0.0348, 0.0693, 0.1037)
selectDetections()
detectionToAnnotationDistances(true)
detectionCentroidDistances(true)
```

Iba1 NVA

```
setImageType('BRIGHTFIELD_H_DAB');
setColorDeconvolutionStains('{"Name" : "H-DAB default", "Stain 1" : "Hematoxylin", "Values 1" :
"0.65111 0.70119 0.29049 ", "Stain 2" : "DAB", "Values 2" : "0.26917 0.56824 0.77759 ",
"Background" : " 255 255 255 "}');
createSelectAllObject(true);
selectAnnotations ();
runPlugin('qupath.imagej.superpixels.DoGSuperpixelsPlugin', '{"downsampleFactor": 1.0,
"sigmaPixels": 8.0,  "minThreshold": 10.0,  "maxThreshold": 230.0,  "noiseThreshold": 1.0}');
selectDetections();
runPlugin('qupath.lib.algorithms.IntensityFeaturesPlugin', '{"downsample": 1.0,  "region": "ROI",
"tileSizePixels": 200.0,  "colorOD": false,  "colorStain1": false,  "colorStain2": true,  "colorStain3": false,
"colorRed": false,  "colorGreen": false,  "colorBlue": false,  "colorHue": false,  "colorSaturation": false,
"colorBrightness": false,  "doMean": true,  "doStdDev": true,  "doMinMax": true,  "doMedian": true,
"doHaralick": false,  "haralickDistance": 1,  "haralickBins": 32}');
addShapeMeasurements("AREA", "LENGTH", "CIRCULARITY", "SOLIDITY", "MAX_DIAMETER",
"MIN_DIAMETER")
setDetectionIntensityClassifications("ROI: 1.00 px per pixel: DAB: Mean", 0.0512, 0.1002, 0.1491)
detectionCentroidDistances(true)
```

Iba1 VA

```
setImageType('BRIGHTFIELD_H_DAB');
setColorDeconvolutionStains('{"Name" : "H-DAB default", "Stain 1" : "Hematoxylin", "Values 1" :
"0.65111 0.70119 0.29049 ", "Stain 2" : "DAB", "Values 2" : "0.26917 0.56824 0.77759 ",
"Background" : " 255 255 255 "}');
selectAnnotations ();
def region = getPathClass('Other')
def other = getPathClass('Region*')
getAnnotationObjects().eachWithIndex { annotation , i ->
  if (i % 2 == 0)
    annotation.setPathClass(region)
 else
    annotation.setPathClass(other)
}
fireHierarchyUpdate()
makeInverseAnnotation()
runPlugin('qupath.imagej.superpixels.DoGSuperpixelsPlugin', '{"downsampleFactor": 1.0,
"sigmaPixels": 8.0,  "minThreshold": 10.0,  "maxThreshold": 230.0,  "noiseThreshold": 1.0}');
selectDetections()
runPlugin('qupath.lib.algorithms.IntensityFeaturesPlugin', '{"downsample": 1.0,  "region": "ROI",
"tileSizePixels": 200.0,  "colorOD": false,  "colorStain1": false,  "colorStain2": true,  "colorStain3": false,
"colorRed": false,  "colorGreen": false,  "colorBlue": false,  "colorHue": false,  "colorSaturation": false,
"colorBrightness": false,  "doMean": true,  "doStdDev": true,  "doMinMax": true,  "doMedian": true,
"doHaralick": false,  "haralickDistance": 1,  "haralickBins": 32}');
```

```
addShapeMeasurements("AREA", "LENGTH", "CIRCULARITY", "SOLIDITY", "MAX_DIAMETER",
"MIN_DIAMETER")
setDetectionIntensityClassifications("ROI: 1.00 px per pixel: DAB: Mean", 0.0512, 0.1002, 0.1491)
selectDetections()
detectionToAnnotationDistances(true)
detectionCentroidDistances(true)
```

GFAP NVA

```
setImageType('BRIGHTFIELD_H_DAB');
setColorDeconvolutionStains('{"Name" : "H-DAB default", "Stain 1" : "Hematoxylin", "Values 1" :
"0.65111 0.70119 0.29049 ", "Stain 2" : "DAB", "Values 2" : "0.26917 0.56824 0.77759 ",
"Background" : " 255 255 255 "}');
createSelectAllObject(true);
selectAnnotations ();
runPlugin('qupath.imagej.superpixels.DoGSuperpixelsPlugin', '{"downsampleFactor": 1.0,
"sigmaPixels": 8.0,  "minThreshold": 10.0,  "maxThreshold": 230.0,  "noiseThreshold": 1.0}');
selectDetections();
runPlugin('qupath.lib.algorithms.IntensityFeaturesPlugin', '{"downsample": 1.0,  "region": "ROI",
"tileSizePixels": 200.0,  "colorOD": false,  "colorStain1": false,  "colorStain2": true,  "colorStain3": false,
"colorRed": false,  "colorGreen": false,  "colorBlue": false,  "colorHue": false,  "colorSaturation": false,
"colorBrightness": false,  "doMean": true,  "doStdDev": true,  "doMinMax": true,  "doMedian": true,
"doHaralick": false,  "haralickDistance": 1,  "haralickBins": 32}');
addShapeMeasurements("AREA", "LENGTH", "CIRCULARITY", "SOLIDITY", "MAX_DIAMETER",
"MIN_DIAMETER")
setDetectionIntensityClassifications("ROI: 1.00 px per pixel: DAB: Mean", 0.1666, 0.2295, 0.2925)
detectionCentroidDistances(true)
```

GFAP VA

```
setImageType('BRIGHTFIELD_H_DAB');
setColorDeconvolutionStains('{"Name" : "H-DAB default", "Stain 1" : "Hematoxylin", "Values 1" :
"0.65111 0.70119 0.29049 ", "Stain 2" : "DAB", "Values 2" : "0.26917 0.56824 0.77759 ",
"Background" : " 255 255 255 "}');
selectAnnotations ();
def region = getPathClass('Other')
def other = getPathClass('Region*')
getAnnotationObjects().eachWithIndex { annotation , i ->
   if (i % 2 == 0)
      annotation.setPathClass(region)
  else
      annotation.setPathClass(other)
}
fireHierarchyUpdate()
makeInverseAnnotation()
runPlugin('qupath.imagej.superpixels.DoGSuperpixelsPlugin', '{"downsampleFactor": 1.0,
"sigmaPixels": 8.0,  "minThreshold": 10.0,  "maxThreshold": 230.0,  "noiseThreshold": 1.0}');
selectDetections()
runPlugin('qupath.lib.algorithms.IntensityFeaturesPlugin', '{"downsample": 1.0,  "region": "ROI",
"tileSizePixels": 200.0,  "colorOD": false,  "colorStain1": false,  "colorStain2": true,  "colorStain3": false,
"colorRed": false,  "colorGreen": false,  "colorBlue": false,  "colorHue": false,  "colorSaturation": false,
"colorBrightness": false,  "doMean": true,  "doStdDev": true,  "doMinMax": true,  "doMedian": true,
"doHaralick": false,  "haralickDistance": 1,  "haralickBins": 32}');
addShapeMeasurements("AREA", "LENGTH", "CIRCULARITY", "SOLIDITY", "MAX_DIAMETER",
"MIN_DIAMETER")
setDetectionIntensityClassifications("ROI: 1.00 px per pixel: DAB: Mean", 0.1666, 0.2295, 0.2925)
selectDetections()
detectionToAnnotationDistances(true)
```

```
detectionCentroidDistances(true)
```

FUS NVA

```
setImageType('BRIGHTFIELD_H_DAB');
setColorDeconvolutionStains('{"Name" : "H-DAB default", "Stain 1" : "Hematoxylin", "Values 1" :
"0.65111 0.70119 0.29049 ", "Stain 2" : "DAB", "Values 2" : "0.26917 0.56824 0.77759 ",
"Background" : " 255 255 255 "}');
createSelectAllObject(true);
selectAnnotations ();
runPlugin('qupath.imagej.superpixels.DoGSuperpixelsPlugin', '{"downsampleFactor": 1.0,
"sigmaPixels": 5.0,  "minThreshold": 10.0,  "maxThreshold": 230.0,  "noiseThreshold": 1.0}');
selectDetections();
runPlugin('qupath.lib.algorithms.IntensityFeaturesPlugin', '{"downsample": 1.0,  "region": "ROI",
"tileSizePixels": 200.0,  "colorOD": false,  "colorStain1": false,  "colorStain2": true,  "colorStain3": false,
"colorRed": false,  "colorGreen": false,  "colorBlue": false,  "colorHue": false,  "colorSaturation": false,
"colorBrightness": false,  "doMean": true,  "doStdDev": true,  "doMinMax": true,  "doMedian": true,
"doHaralick": false,  "haralickDistance": 1,  "haralickBins": 32}');
addShapeMeasurements("AREA", "LENGTH", "CIRCULARITY", "SOLIDITY", "MAX_DIAMETER",
"MIN_DIAMETER")
setDetectionIntensityClassifications("ROI: 1.00 px per pixel: DAB: Mean", 0.0900, 0.1100, 0.1300)
detectionCentroidDistances(true)
```

FUS VA

```
setImageType('BRIGHTFIELD_H_DAB');
setColorDeconvolutionStains('{"Name" : "H-DAB default", "Stain 1" : "Hematoxylin", "Values 1" :
"0.65111 0.70119 0.29049 ", "Stain 2" : "DAB", "Values 2" : "0.26917 0.56824 0.77759 ",
"Background" : " 255 255 255 "}');
selectAnnotations ();
def region = getPathClass('Other')
def other = getPathClass('Region*')
getAnnotationObjects().eachWithIndex { annotation , i ->
  if (i % 2 == 0)
     annotation.setPathClass(region)
 else
     annotation.setPathClass(other)
}
fireHierarchyUpdate()
makeInverseAnnotation()
runPlugin('qupath.imagej.superpixels.DoGSuperpixelsPlugin', '{"downsampleFactor": 1.0,
"sigmaPixels": 5.0,  "minThreshold": 10.0,  "maxThreshold": 230.0,  "noiseThreshold": 1.0}');
selectDetections()
runPlugin('qupath.lib.algorithms.IntensityFeaturesPlugin', '{"downsample": 1.0,  "region": "ROI",
"tileSizePixels": 200.0,  "colorOD": false,  "colorStain1": false,  "colorStain2": true,  "colorStain3": false,
"colorRed": false,  "colorGreen": false,  "colorBlue": false,  "colorHue": false,  "colorSaturation": false,
"colorBrightness": false,  "doMean": true,  "doStdDev": true,  "doMinMax": true,  "doMedian": true,
"doHaralick": false,  "haralickDistance": 1,  "haralickBins": 32}');
addShapeMeasurements("AREA", "LENGTH", "CIRCULARITY", "SOLIDITY", "MAX_DIAMETER",
"MIN_DIAMETER")
setDetectionIntensityClassifications("ROI: 1.00 px per pixel: DAB: Mean", 0.0900, 0.1100, 0.1300)
selectDetections()
detectionToAnnotationDistances(true)
detectionCentroidDistances(true)
```

TDP43 NVA

```
setImageType('BRIGHTFIELD_H_DAB');
setColorDeconvolutionStains('{"Name" : "H-DAB default", "Stain 1" : "Hematoxylin", "Values 1" :
"0.65111 0.70119 0.29049 ", "Stain 2" : "DAB", "Values 2" : "0.26917 0.56824 0.77759 ",
"Background" : " 255 255 255 "}');
createSelectAllObject(true);
selectAnnotations ();
runPlugin('qupath.imagej.superpixels.DoGSuperpixelsPlugin', '{"downsampleFactor": 1.0,
"sigmaPixels": 5.0,  "minThreshold": 10.0,  "maxThreshold": 230.0,  "noiseThreshold": 1.0}');
selectDetections();
runPlugin('qupath.lib.algorithms.IntensityFeaturesPlugin', '{"downsample": 1.0,  "region": "ROI",
"tileSizePixels": 200.0,  "colorOD": false,  "colorStain1": false,  "colorStain2": true,  "colorStain3": false,
"colorRed": false,  "colorGreen": false,  "colorBlue": false,  "colorHue": false,  "colorSaturation": false,
"colorBrightness": false,  "doMean": true,  "doStdDev": true,  "doMinMax": true,  "doMedian": true,
"doHaralick": false,  "haralickDistance": 1,  "haralickBins": 32}');
addShapeMeasurements("AREA", "LENGTH", "CIRCULARITY", "SOLIDITY", "MAX_DIAMETER",
"MIN_DIAMETER")
setDetectionIntensityClassifications("ROI: 1.00 px per pixel: DAB: Mean", 0.0719, 0.1223, 0.1713)
detectionCentroidDistances(true)
```

TDP43 VA

```
setImageType('BRIGHTFIELD_H_DAB');
setColorDeconvolutionStains('{"Name" : "H-DAB default", "Stain 1" : "Hematoxylin", "Values 1" :
"0.65111 0.70119 0.29049 ", "Stain 2" : "DAB", "Values 2" : "0.26917 0.56824 0.77759 ",
"Background" : " 255 255 255 "}');
selectAnnotations ();
def region = getPathClass('Other')
def other = getPathClass('Region*')
getAnnotationObjects().eachWithIndex { annotation , i ->
  if (i % 2 == 0)
    annotation.setPathClass(region)
  else
    annotation.setPathClass(other)
}
fireHierarchyUpdate()
makeInverseAnnotation()
runPlugin('qupath.imagej.superpixels.DoGSuperpixelsPlugin', '{"downsampleFactor": 1.0,
"sigmaPixels": 5.0,  "minThreshold": 10.0,  "maxThreshold": 230.0,  "noiseThreshold": 1.0}');
selectDetections()
runPlugin('qupath.lib.algorithms.IntensityFeaturesPlugin', '{"downsample": 1.0,  "region": "ROI",
"tileSizePixels": 200.0,  "colorOD": false,  "colorStain1": false,  "colorStain2": true,  "colorStain3": false,
"colorRed": false,  "colorGreen": false,  "colorBlue": false,  "colorHue": false,  "colorSaturation": false,
"colorBrightness": false,  "doMean": true,  "doStdDev": true,  "doMinMax": true,  "doMedian": true,
"doHaralick": false,  "haralickDistance": 1,  "haralickBins": 32}');
addShapeMeasurements("AREA", "LENGTH", "CIRCULARITY", "SOLIDITY", "MAX_DIAMETER",
"MIN_DIAMETER")
setDetectionIntensityClassifications("ROI: 1.00 px per pixel: DAB: Mean", 0.0719, 0.1223, 0.1713)
selectDetections()
detectionToAnnotationDistances(true)
detectionCentroidDistances(true)
```

FUS NVA neurons localisation

```
setImageType('BRIGHTFIELD_H_DAB');
```

```
setColorDeconvolutionStains('{"Name" : "H-DAB default", "Stain 1" : "Hematoxylin", "Values 1" : "0.65111 0.70119 0.29049 ", "Stain 2" : "DAB", "Values 2" : "0.26917 0.56824 0.77759 ", "Background" : " 255 255 255 "}');
setPixelSizeMicrons(0.625,0.625)

createSelectAllObject(true);
selectAnnotations ();

runPlugin('qupath.imagej.detect.cells.WatershedCellDetection', '{"detectionImageBrightfield": "Hematoxylin OD",  "requestedPixelSizeMicrons": 0.5,  "backgroundRadiusMicrons": 0.0,  "medianRadiusMicrons": 0.0,  "sigmaMicrons": 4.0,  "minAreaMicrons": 150.0,  "maxAreaMicrons": 5000.0,  "threshold": 0.1,  "maxBackground": 2.0,  "watershedPostProcess": true,  "excludeDAB": false,  "cellExpansionMicrons": 10.0,  "includeNuclei": true,  "smoothBoundaries": true,  "makeMeasurements": true}');
```

FUS VA neurons localisation

```
setImageType('BRIGHTFIELD_H_DAB');
setColorDeconvolutionStains('{"Name" : "H-DAB default", "Stain 1" : "Hematoxylin", "Values 1" : "0.65111 0.70119 0.29049 ", "Stain 2" : "DAB", "Values 2" : "0.26917 0.56824 0.77759 ", "Background" : " 255 255 255 "}');
setPixelSizeMicrons(0.625,0.625)

selectAnnotations ();
def region = getPathClass('Other')
def other = getPathClass('Region*')
getAnnotationObjects().eachWithIndex { annotation , i ->
   if (i % 2 == 0)
      annotation.setPathClass(region)
  else
      annotation.setPathClass(other)
}
fireHierarchyUpdate()
makeInverseAnnotation()


runPlugin('qupath.imagej.detect.cells.WatershedCellDetection', '{"detectionImageBrightfield": "Hematoxylin OD",  "requestedPixelSizeMicrons": 0.5,  "backgroundRadiusMicrons": 0.0,  "medianRadiusMicrons": 0.0,  "sigmaMicrons": 4.0,  "minAreaMicrons": 150.0,  "maxAreaMicrons": 5000.0,  "threshold": 0.1,  "maxBackground": 2.0,  "watershedPostProcess": true,  "excludeDAB": false,  "cellExpansionMicrons": 10.0,  "includeNuclei": true,  "smoothBoundaries": true,  "makeMeasurements": true}');
```

FUS VA glial localisation

```
setImageType('BRIGHTFIELD_H_DAB');
setColorDeconvolutionStains('{"Name" : "H-DAB default", "Stain 1" : "Hematoxylin", "Values 1" : "0.65111 0.70119 0.29049 ", "Stain 2" : "DAB", "Values 2" : "0.26917 0.56824 0.77759 ", "Background" : " 255 255 255 "}');
setPixelSizeMicrons(0.625,0.625)

selectAnnotations ();
def region = getPathClass('Other')
def other = getPathClass('Region*')
getAnnotationObjects().eachWithIndex { annotation , i ->
   if (i % 2 == 0)
      annotation.setPathClass(region)
  else
```

```
    annotation.setPathClass(other)
}
fireHierarchyUpdate()
makeInverseAnnotation()


runPlugin('qupath.imagej.detect.cells.WatershedCellDetection', '{"detectionImageBrightfield": "Hematoxylin OD",  "requestedPixelSizeMicrons": 0.5,  "backgroundRadiusMicrons": 0.0,  "medianRadiusMicrons": 0.0,  "sigmaMicrons": 4.0,  "minAreaMicrons": 10.0,  "maxAreaMicrons": 100.0,  "threshold": 0.1,  "maxBackground": 2.0,  "watershedPostProcess": true,  "excludeDAB": false,  "cellExpansionMicrons": 5.0,  "includeNuclei": true,  "smoothBoundaries": true,  "makeMeasurements": true}');
```

FUS NVA glial localisation

```
setImageType('BRIGHTFIELD_H_DAB');
setColorDeconvolutionStains('{"Name" : "H-DAB default", "Stain 1" : "Hematoxylin", "Values 1" : "0.65111 0.70119 0.29049 ", "Stain 2" : "DAB", "Values 2" : "0.26917 0.56824 0.77759 ", "Background" : " 255 255 255 "}');
setPixelSizeMicrons(0.625,0.625)

createSelectAllObject(true);
selectAnnotations ();

runPlugin('qupath.imagej.detect.cells.WatershedCellDetection', '{"detectionImageBrightfield": "Hematoxylin OD",  "requestedPixelSizeMicrons": 0.5,  "backgroundRadiusMicrons": 0.0,  "medianRadiusMicrons": 0.0,  "sigmaMicrons": 4.0,  "minAreaMicrons": 10.0,  "maxAreaMicrons": 100.0,  "threshold": 0.1,  "maxBackground": 2.0,  "watershedPostProcess": true,  "excludeDAB": false,  "cellExpansionMicrons": 5.0,  "includeNuclei": true,  "smoothBoundaries": true,  "makeMeasurements": true}');
```
